# Numerical study of spatial and temporal dynamics of integrin clustering during early cell adhesion

**DOI:** 10.64898/2026.06.07.730653

**Authors:** Kohsuke Tsukui, Tatsuya Kawai, Hiromi Miyoshi, Naoya Sakamoto, Hiro Wakimura, Satoshi Ii

## Abstract

Integrins are adhesion proteins that diffuse along the cell membrane, bind extracellular ligands, and form clusters during early cell adhesion. Although integrin clustering and its spatial organization are important for subsequent biological processes, the mechanisms that generate cellular-scale patterns remain unclear. To investigate these mechanisms, we developed a multiscale cell adhesion model coupling membrane mechanics with integrin dynamics. Membrane deformation was represented using a hybrid continuous/discrete model, whereas integrin motion and binding kinetics were described using Brownian dynamics combined with a transition-state model. Simulated bound-integrin distributions were converted into pseudo-fluorescence images and compared with experimental paxillin fluorescence images. Most experimental and simulated images exhibited a peripheral bias, characterized by higher peripheral than central intensity, and comparison of metrics based on radial intensity distributions and image-derived clusters indicated that the simulations reproduced part of the experimental variation. Without the integrin–ligand bond force acting on the membrane, bound integrins did not form clearly localized high-density clusters, although a weak peripheral bias persisted; with the force, local membrane deformation promoted the formation of localized high-density clusters. Increasing the spreading speed shifted integrin clustering from a smaller number of large clusters to a larger number of small clusters and altered their spatial organization. These findings indicate that the spatial organization of integrin clusters depends on both binding-induced membrane deformation and the relative timescales of membrane spreading and integrin binding.

## 1 Introduction

Cell adhesion involves cellular-scale membrane deformation and molecular-scale interactions mediated by transmembrane proteins and is essential for various biological functions, including migration, division, and cell differentiation (Shinde et al., 2021; Ferrai and Schulte, 2024). In the initial stage of cell adhesion, membrane attachment is induced by physical adsorption originating from electro-chemical interactions between the cell membrane and the substrate surface. Among the involved components are integrins, transmembrane adhesion proteins that diffuse along the cell membrane and bind to ligand molecules, such as fibronectin and collagen, on the substrate surface. Subsequently, other integrins and intracellular adhesion-related proteins, such as paxillin and talin, are recruited around the bound integrins, resulting in the formation of integrin clusters. These clusters form the core of nascent adhesions (NAs).

Integrin clustering, followed by NA formation, and the spatial distribution of these clusters are important for understanding and controlling cell dynamics. Integrin clusters serve as initial sites for the recruitment of adhesion-related proteins that subsequently form mature focal adhesions (FAs), and they are also involved in early signaling and mechanosensing (Henning Stumpf et al., 2020). In addition, integrin-based adhesion structures are not static assemblies but dynamic structures that are reorganized in response to the local status of the extracellular matrix, the cytoskeleton, and signaling pathways mediated by adhesion-related proteins (Wolfenson et al., 2013, 2014). Integrin clustering and FA maturation are promoted by traction forces generated through coupling with the cytoskeleton and by mechanotransductive signaling associated with talin extension (Balaban et al., 2001; Pernier et al., 2023). Even when the size of mature FAs remains unchanged, FAs located in the periphery of the adhesion region can transmit higher traction forces than those located closer to the cell center (Stricker et al., 2011). Thus, the formation dynamics and spatial distribution of integrin clusters across the cell during the early stage of adhesion are important, including for subsequent maturation into FAs. In particular, positional differences within the region in which a cell adheres to a substrate, such as between its periphery and its center, may influence traction-force transmission and FA maturation.

Experimental approaches have revealed the size and mean lifetime of NAs, as well as the fact that NAs form prior to substrate-rigidity sensing and the associated generation of cellular contractile forces (Changede et al., 2015). In addition, live-cell imaging of paxillin, an adhesion-associated protein recruited to NAs, has shown that paxillin-positive structures are enriched near the edge of the cell–substrate adhesion region during the early stages of cell spreading (Fouchard et al., 2014). Transient NAs have also been visualized during the initial stage of cell adhesion (Lee et al., 2022). However, because of the low signal-to-noise ratio, it remains difficult to directly measure the binding dynamics of individual integrins at the whole-cell scale. Therefore, it remains unclear how membrane–substrate interactions and integrin dynamics collectively shape the cellular-scale spatial organization of early adhesion-associated structures.

Numerical simulations are a promising approach to elucidate the mechanisms of integrin clustering. These include microscopic models that focus on the molecular dynamics of integrin–ligand interactions (Tong et al., 2023), mesoscopic models that describe interactions between the cell membrane and the substrate surface (Paszek et al., 2009; Bidone et al., 2019; Carney et al., 2023; Liang et al., 2024; Blanchard, 2024), and macroscopic models that link cell adhesion and deformation to receptor–ligand interaction (Kim et al., 2013; Golestaneh and Nadler, 2016; Vargas et al., 2020). Microscopic models can resolve protein-level conformational changes and the molecular mechanisms of integrins. However, the time scales for which such simulations can be generated are limited, making it difficult to capture adhesion-site dynamics in which multiple integrins cluster and associated proteins are recruited and interact. In mesoscopic models, integrin motion is typically represented as particle diffusion, and integrin–ligand bond formation is modeled as a stochastic process. A recent model represented integrins using active surface density and a probability distribution for the position of a ligand-paired integrin under steady-state approximations (Blanchard, 2024). These models have clarified how substrate stiffness, glycocalyx and cytoskeletal structures influence integrin clustering. Nevertheless, because of their limited spatial scale and the frequent use of periodic boundary conditions, they are not well suited to investigating cluster distributions across an entire cell. In addition, they mainly target already-spread adherent cells and therefore cannot describe integrin clustering during membrane adhesion and spreading. Macroscopic models are suitable for analyzing cellular-scale mechanical behavior and successfully describe receptor–ligand-mediated cell adhesion and deformation, as well as cell migration driven by traction forces generated by FAs and the associated cytoskeleton. However, the lateral transport and clustering of individual integrins on the membrane are not explicitly represented. Moreover, FA formation is simplified as bond formation between fixed points on the membrane and the substrate. Therefore, these models cannot describe the process of formation of adhesion sites or local mechanics at the membrane surface.

To bridge the gap between these spatial scales and investigate integrin clustering during early cell adhesion, we propose a multiscale mechanical model that couples whole-cell membrane mechanics with mesoscopic integrin dynamics. The cell membrane dynamics was modeled by conceptualizing the cell as a capsule made up of triangular mesh elements within a hybrid continuous/discrete mechanics framework. The integrin dynamics was modeled mesoscopically by representing each integrin as a particle undergoing Brownian dynamics with a transition-state model for bond formation. The cell membrane dynamics and the integrin dynamics were mechanically coupled by integrating information on the force of the binding between integrins and their ligands. We simulated integrin clustering on the adhering membrane and analyzed the spatiotemporal distribution of integrin clusters at the cell scale. Simulation results were converted into pseudo-fluorescence images using a point spread function, and their spatial patterns were compared with those observed in experimental paxillin fluorescence images during the early stages of adhesion. We examined the time evolution of cell adhesion and integrin clustering and evaluated how membrane stiffness, the contribution of the integrin–ligand bond force, and membrane spreading speed affect cluster dynamics and spatial distributions.

## 2 Methods

### 2.1 Cell adhesion mechanical model

Our multiscale model incorporated both the macroscopic dynamics of cell membrane deformation and the mesoscopic dynamics of integrin along the cell membrane (Fig. 1a). The cell was modeled as a capsule made up of triangular mesh elements, and integrin was modeled as particles undergoing two-dimensional diffusion along each surface mesh element (Fig. 1b). When the displacement indicated that the particle had left the current element, the excess displacement was rotated about the shared edge and mapped onto the neighboring element (Fig. 1c).

**Figure 1:**
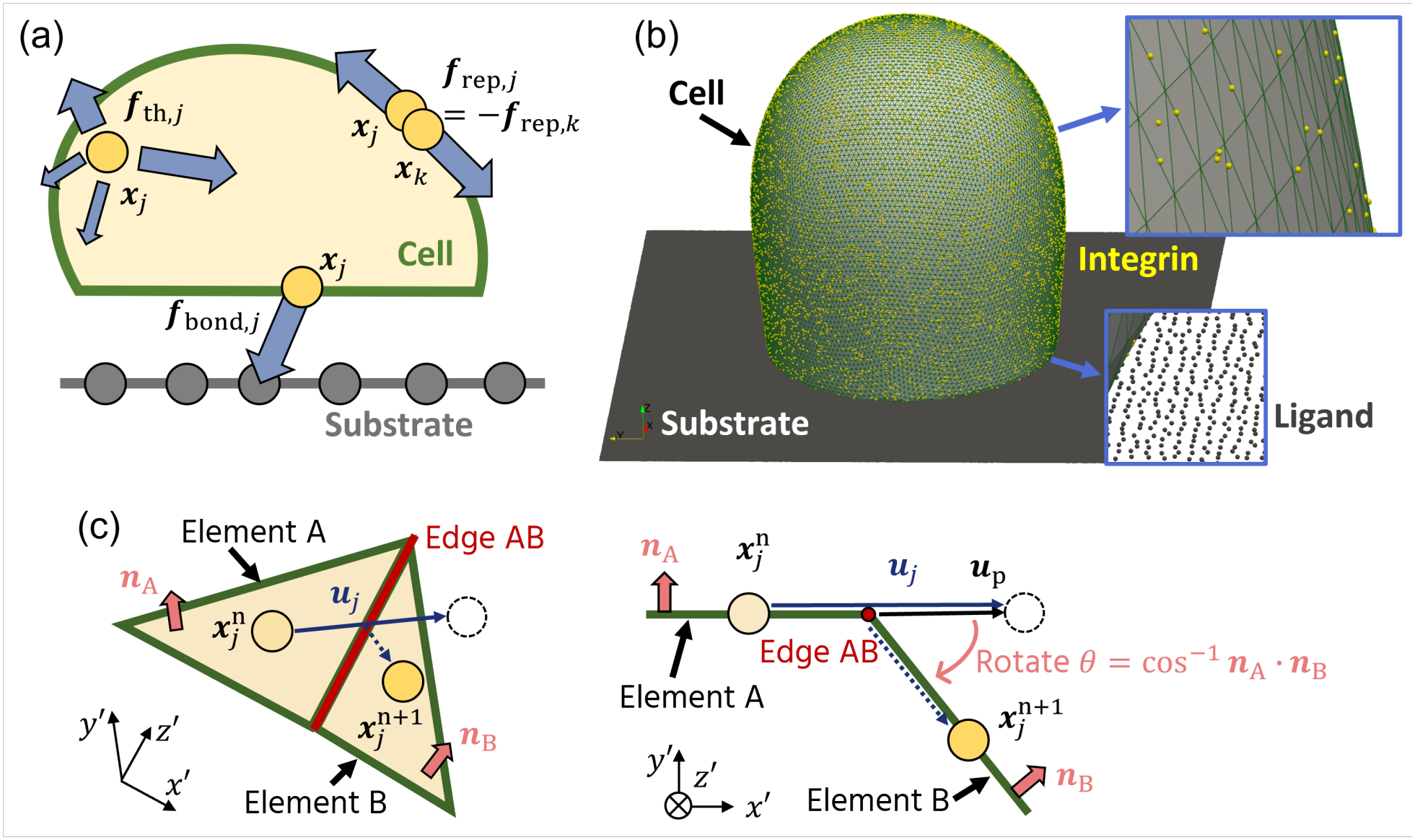
Overview of multiscale cell adhesion model. **a** Integrins move on the adhesive cell membrane and can bind to ligands on the substrate. **b** The cell membrane, the integrin, and the ligand were modeled as a capsule made up of triangular mesh elements, a particle on the element, and a particle confined to the substrate surface, respectively. **c** Movement of the integrin to a neighboring element. Integrins were modeled as particles undergoing two-dimensional diffusion on membrane elements. When the displacement indicated that the particle had left the current element (Element A), the excess displacement **u**_p_ was rotated about the shared edge (Edge AB) and mapped onto the neighboring element (Element B).

#### 2.1.1 Modeling of integrin dynamics

Integrins were modeled as particles undergoing two-dimensional diffusion on each membrane element. Integrin movement was described using a Langevin equation and given as

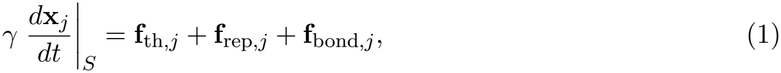

where

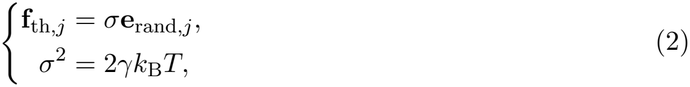

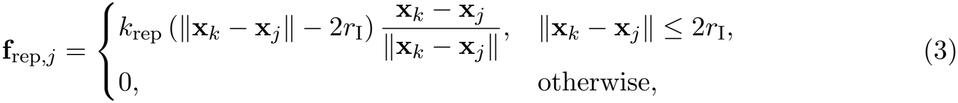

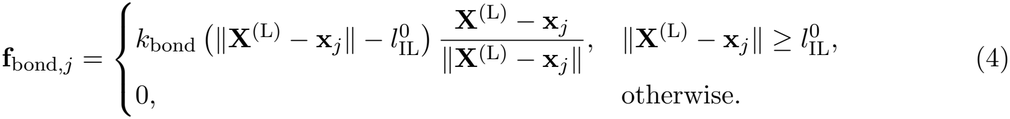

Here, **x**_*j*_ is the position of the *j*-th integrin node, *γ* is the friction constant, **f**_th,*j*_ is the thermal force, *σ* is the random force magnitude, *k*_B_ is the Boltzmann constant, *T* is the absolute temperature, **e**_rand,*j*_ is the random vector, **f**_rep,*j*_ is the repulsive force between integrins, *k*_rep_ is the repulsive constant, *r*_I_ is the radius of integrin, **f**_bond,*j*_ is the bond force between the integrin and the ligand on the substrate, 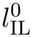 is the natural length of the integrin–ligand bond, and **X**^(L)^ is the position of the ligand node bound to the *j*-th integrin. The bond force was applied only to integrins bound to ligands; the corresponding binding states are described later. Here, *d/dt*|_*S*_ denotes the time derivative in a frame moving with the membrane surface, meaning that integrin nodes move together with the membrane as it undergoes local motion. The friction coefficient *γ* was determined from the Einstein relation as

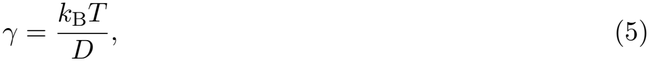

where *D* is an effective lateral diffusion coefficient determined from experimentally reported integrin mobility (Table 1). By discretizing Eq. (1) (Ermak and McCammon, 1978), the thermal force in Eq. (2) became

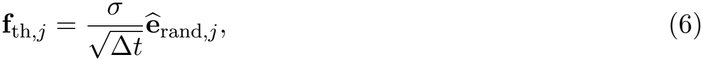

where Δ*t* is the time interval and 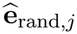 is the random number satisfying the standard normal distribution.

**Table 1:**
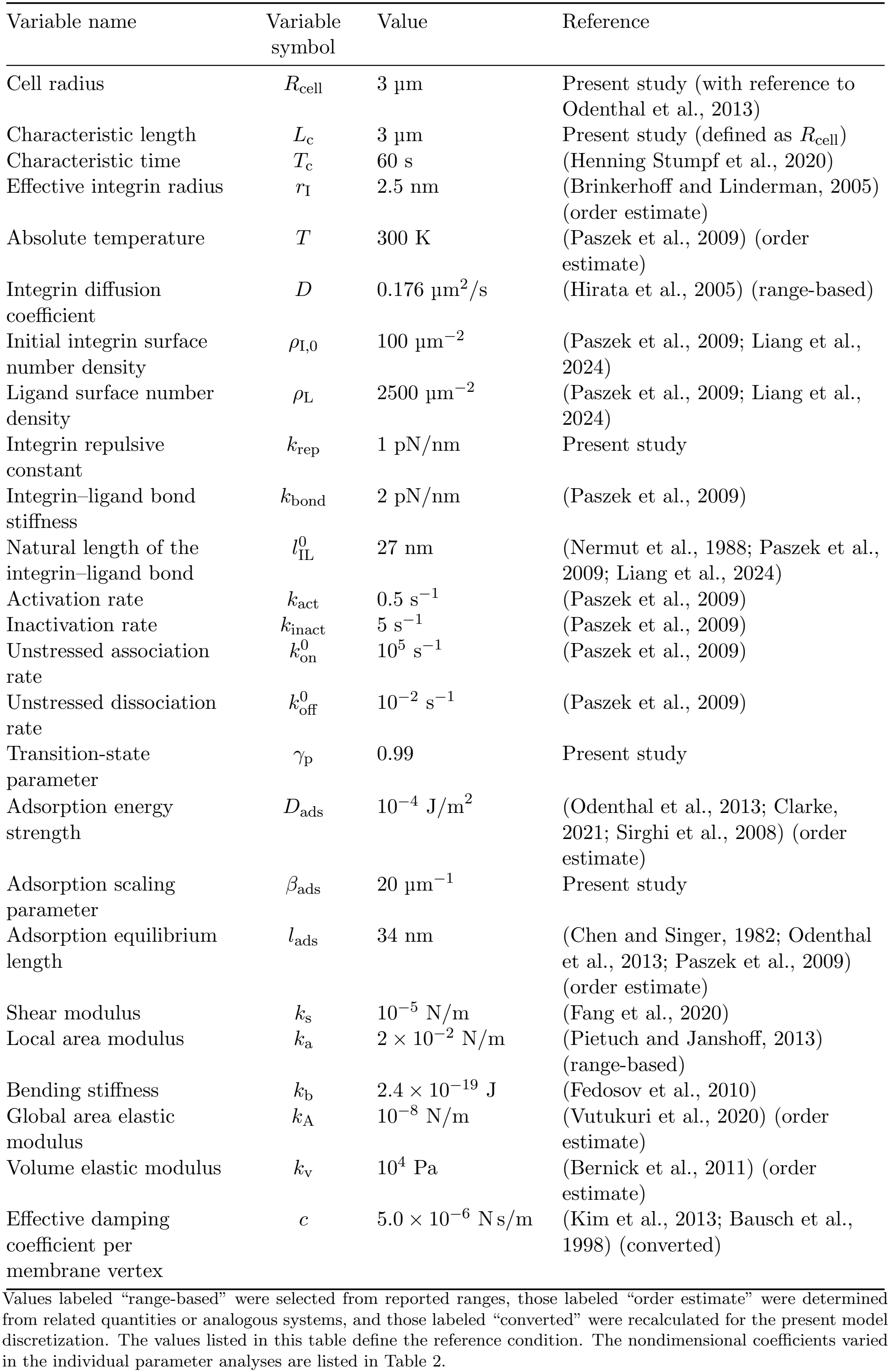
Physical parameters used in the simulations.

Depending on the underlying molecular mechanisms, integrins can adopt one of three states: inactive, active, or bound. In the active state, the integrin can bind to the ligand. In our model, the integrin activation and binding kinetics with the ligand were modeled as a stochastic process using a kinetic Monte Carlo method (Kim et al., 2013; Ii et al., 2018). Each probability and reaction rate were summarized as follows:

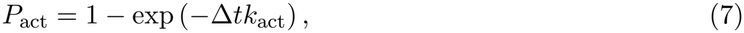

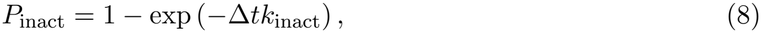

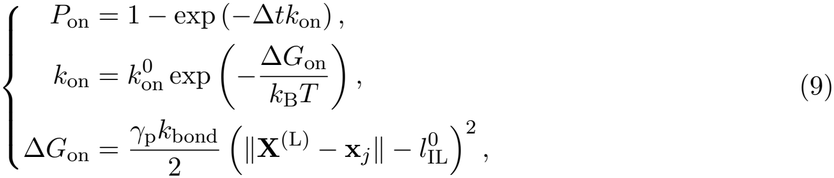

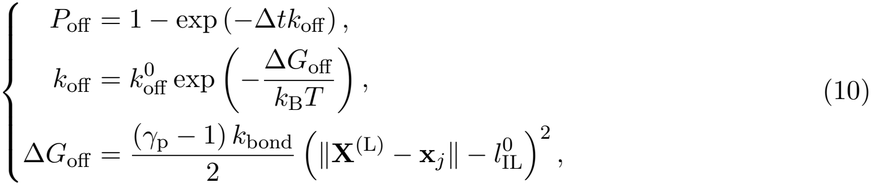

where *P*_*α*_ (*α* = act, inact, on, off) is the probability of changing to state *α, k*_*α*_ is the reaction rate for state *α*, 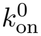 and 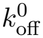 are the unstressed reaction rates, and *γ*_p_ is the transition-state parameter. In the bond formation dynamics of the *j*-th activated integrin, the ligand node closest to the *j*-th integrin is determined. Ligand nodes were placed on a square lattice on the substrate surface, with each node independently displaced from the center of its lattice cell by a uniformly distributed random offset of up to one-quarter of the lattice spacing in each in-plane direction.

#### 2.1.2 Modeling of cell membrane

The cell membrane was modeled as a hyperelastic membrane made up of triangular mesh elements. The mechanical equation of the *i*-th vertex of the membrane surface mesh was given as

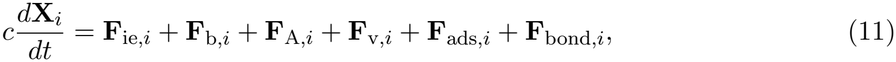

where **X**_*i*_ is the position of the *i*-th vertex, *c* is the effective nodal damping coefficient representing the dissipative coupling of the membrane–cortex complex with the intracellular and extracellular environments, **F**_ie,*i*_ is the in-plane and local area elastic force, **F**_b,*i*_ is the bending force, **F**_A,*i*_ is the global area elastic force, **F**_v,*i*_ is the volume elastic force, **F**_ads,*i*_ is the physical adsorption force caused by interaction between the cell membrane and the substrate, and **F**_bond,*i*_ is the bond force between the integrin and the ligand on the substrate.

The in-plane elastic force per unit area **q** employs the compressive neo-Hookean model and strain energy function *W*_I_ was given as

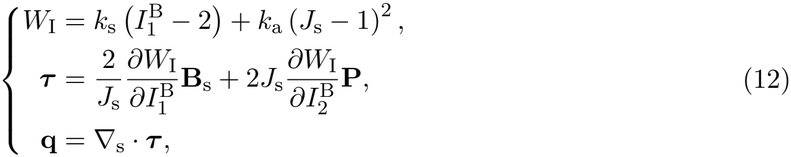

where *k*_s_ is the shear modulus, *k*_a_ is the local area modulus, 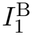 and 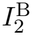 are the first and second invariants of the surface left Cauchy–Green deformation tensor **B**_s_, *J*_s_ is the surface Jacobian, ***τ*** is the membrane tension tensor (Cauchy stress tensor), **F**_s_ is the surface deformation tensor, **P** = **I** − **nn** is the surface projection tensor, **I** is the identity tensor, **n** is the unit normal vector of the membrane, and ∇_s_ = **P**· ∇_*x*_ is the surface gradient operator. In this model, **q** was approximated by the finite element formulation using a linear shape function *N*_*i*_ and **F**_ie,*i*_ was given using internal force at a vertex **q**_*i*_ as

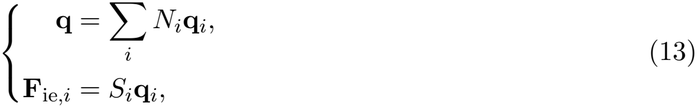

where *S*_*i*_ is the local area associated with the *i*-th vertex, defined as the area of the Voronoi region corresponding to the *i*-th vertex.

The bending energy *E*_b_ was modeled using Helfrich’s energy, and the energy and force were modeled as

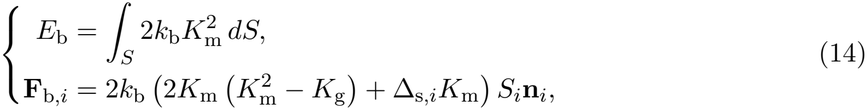

where *k*_b_ is the bending stiffness, *K*_m_ is the mean curvature, *K*_g_ is the Gaussian curvature, Δ_s,*i*_ is the Laplace–Beltrami operator evaluated at the *i*-th vertex, and **n**_*i*_ is the unit normal vector at the *i*-th vertex.

The global area elastic energy and force (Fedosov et al., 2010) were modeled as

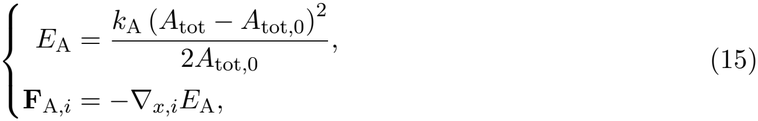

where *k*_A_ is the global area elastic modulus, and *A*_tot_ and *A*_tot,0_ are the surface areas of the cell membrane in the current state and initial state, respectively.

The volume elastic energy and force (Fedosov et al., 2010) were modeled as

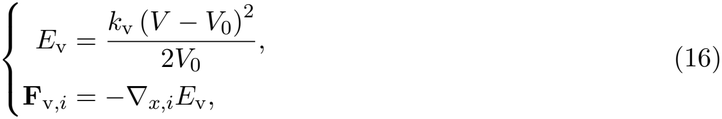

where *k*_v_ is the volume elastic modulus, and *V* and *V*_0_ are the volumes of the cell membrane in the current state and initial state, respectively.

The physical adsorption energy based on the interaction between the cell membrane surface and substrate surface was modeled using the Morse potential as

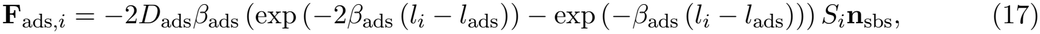

where *D*_ads_ is the energy strength (well depth of potential), *β*_ads_ is the scaling parameter, *l*_*i*_ is the distance between a cell membrane vertex and substrate surface, *l*_ads_ is the equilibrium length, and **n**_sbs_ is the unit normal vector of the substrate surface. In this study, the substrate surface was defined as a plane perpendicular to the *z*-axis, and the substrate normal vector was set to **n**_sbs_ = (0, 0, 1).

The bond force **F**_bond,*i*_ acting on the membrane surface has a role in mechanically coupling with the cell membrane and mesoscopic integrin dynamics and was computed by

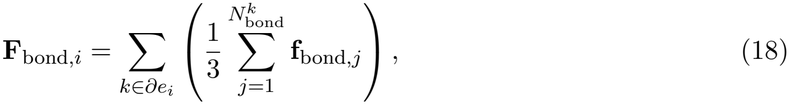

where *∂e*_*i*_ is the set of mesh elements containing the *i*-th vertex, and 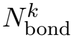 is the number of bound integrins on the *k*-th mesh element. If the mesh resolution is sufficiently fine, the forces acting on each node of the membrane mesh would be approximated by equally distributing the resultant bond force.

#### 2.1.3 Nondimensionalization of the governing equation of the cell membrane

In our model, the physical adsorption energy drives cell adhesion phenomena. For clarity, we derive the dimensionless equations from Eq. (11). Physical parameters were nondimensionalized using the characteristic length *L*_c_, the characteristic time *T*_c_, and the damping constant *c*. The dimensionless equations normalized by the physical adsorption strength *D*_ads_*β*_ads_*L*_c_^2^ were given as

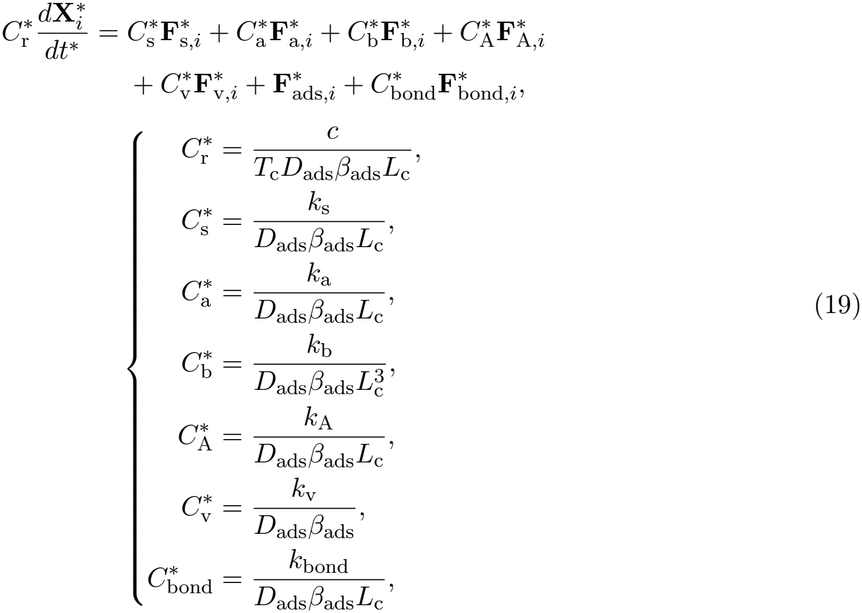

where variables marked with an asterisk * represent dimensionless quantities, 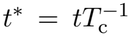 is the normalized time, 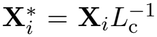 is the normalized position of the *i*-th membrane vertex, 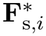 is the in-plane shear elastic force, 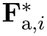 is the in-plane local area elastic force, 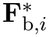 is the bending force, 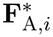 is the global area elastic force, 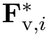 the volume elastic force, 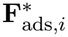 is the physical adsorption force, 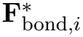 is the bond force, and 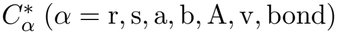 represents the relative contribution of the *α* force to the physical adsorption force. The characteristic length *L*_c_ was defined as the cell radius *R*_cell_ in this study. The characteristic time *T*_c_ was defined based on the average lifetime of an NA (Henning Stumpf et al., 2020) and was set to *T*_c_ = 60 s in this study.

### 2.2 Simulation setup

Although the governing equations were nondimensionalized, representing a cell with the experimentally measured radius while maintaining the prescribed spatial resolution and molecular surface densities would require substantially larger numbers of membrane elements, integrins, and ligands. Therefore, to keep the computational cost tractable, a reduced model cell was used, and its radius was set to *R*_cell_ = 3 µm with reference to the radius of 3.25 µm adopted in a previous deformable-cell spreading model (Odenthal et al., 2013). The total numbers of mesh elements *N*_e_ and vertices *N*_v_ were 30,420 and 15,212, respectively. This mesh resolution was adopted after a mesh-dependence analysis confirmed convergence of the normalized adhesion area and bound-integrin fraction (Fig. S1). The substrate was modeled as a planar surface located at *z* = 0. The cell was initially assumed to be spherical and positioned at a distance from the substrate corresponding to the equilibrium distance of the Morse potential *l*_ads_. On the substrate surface (*z* = 0), ligand nodes were placed within a square region with side length *L*_region_ = 10 µm using the randomly perturbed square lattice described above. Based on the literature values listed in Table 1, the ligand surface number density was prescribed as *ρ*_L_ = 2500 µm^−2^, and the total number of ligand nodes was accordingly set to 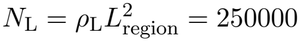. The integrins were initially distributed uniformly with a surface number density of *ρ*_I,0_ = 100 µm^−2^, and the fixed total number of integrins was *N*_I_ = 11300, which gives 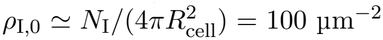. Their initial positions were assigned randomly by selecting mesh elements at random, with integrins distributed randomly on those mesh surfaces.

The physical parameters defining the reference condition are summarized in Table 1, and the corresponding nondimensional coefficients are listed in Table 2.

**Table 2:**
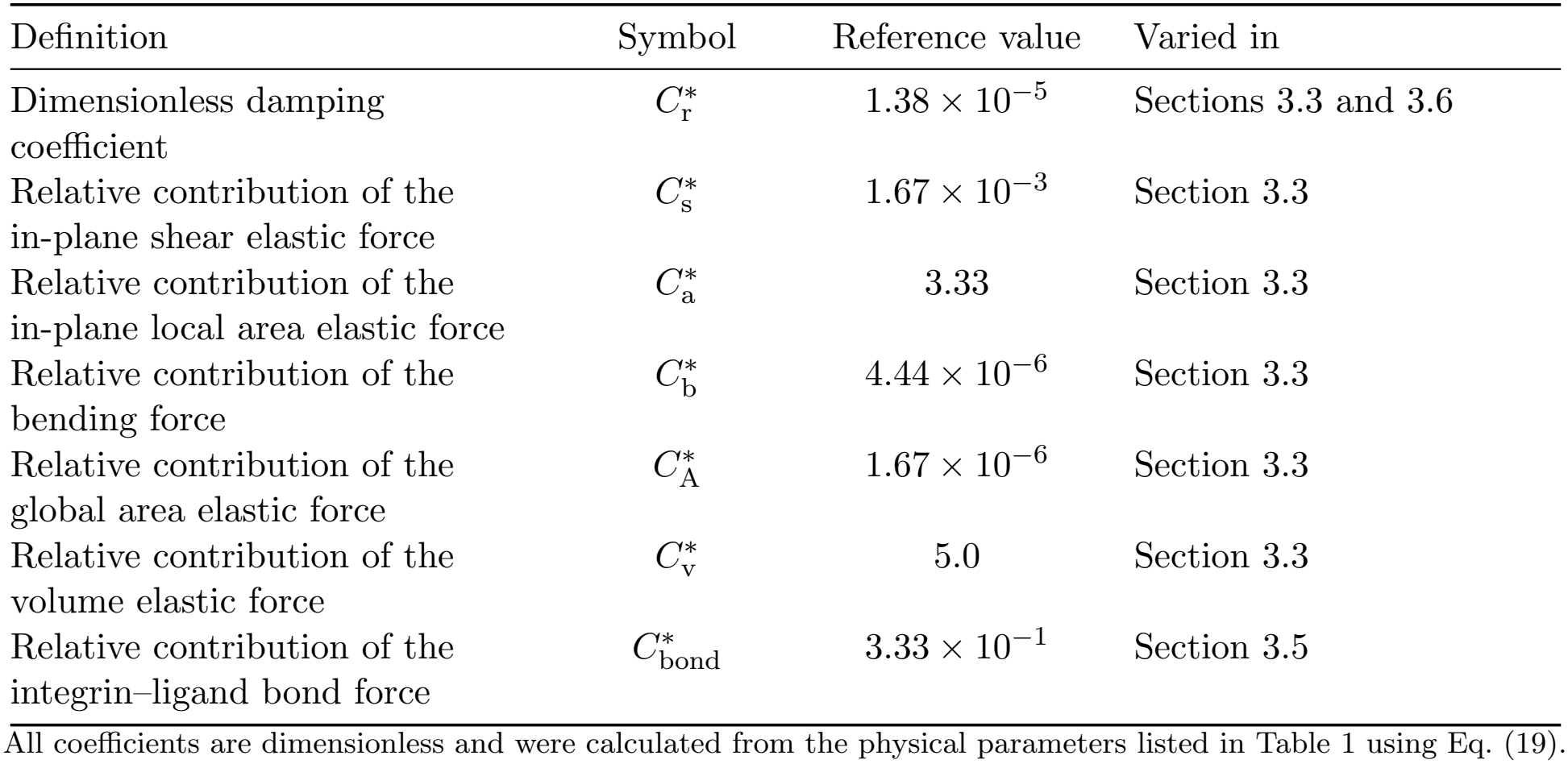
Nondimensional coefficients for the reference condition and the Results subsections in which they were varied.

To account for differences in membrane discretization and mesh resolution, the nodal damping coefficient *c* was converted to an effective damping coefficient per unit membrane area, defined as 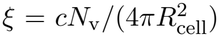, where *N*_v_ is the number of membrane vertices. The values used by Kim et al. (2013), *c* = 1.0×10^−3^ N s*/*m, *N*_v_ = 549, and *R*_cell_ = 8 µm, give *ξ* = 6.83×10^8^ N s*/*m^3^. Applying this value to the present discretization, with *N*_v_ = 15212 and *R*_cell_ = 3 µm, gives 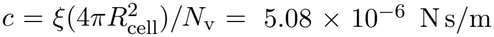. We therefore adopted *c* = 5.0 × 10^−6^ N s*/*m as the reference value. The corresponding value, *ξ* = 6.73 × 10^8^ N s*/*m^3^, is of the same order of magnitude as 2 × 10^9^ N s*/*m^3^, estimated by integrin-coupled magnetic-bead microrheometry in NIH 3T3 fibroblasts (Bausch et al., 1998).

In the parameter analyses presented in the Results, *λ* denotes a common scaling factor applied to the membrane-stiffness-related coefficients 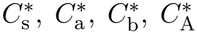 and 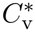 relative to their reference values listed in Table 2. Thus, *λ* = 1 indicates that these coefficients take their reference values. In each Results subsection, only coefficients that differ from the reference condition are specified; unless otherwise stated, all others retain their reference values.

### 2.3 Experiments

To compare the model predictions with experimental observations, we conducted experiments to characterize cell morphology and paxillin-positive clusters during the early stages of adhesion. Mouse fibroblasts (Swiss 3T3 Albino) cultured in Dulbecco’s Modified Eagle’s Medium (Sigma) were used for the experiments. A 500 µL aliquot of a cell suspension at a density of 100 cells/µL was seeded onto 3.5 cm glass-bottom dishes coated with fibronectin (0.02 µg/µL). Each dish was incubated for a specified duration *T*_inc_ (2, 5, and 15 min).

Subsequently, without aspirating the culture medium, 500 µL of 8% paraformaldehyde was added directly to each dish, and the cells were fixed for 15 min (equivalent to fixation with 4% paraformaldehyde). The actin cytoskeleton was stained with phalloidin, and paxillin was stained using rabbit primary and anti-rabbit secondary antibodies. Cell morphology and protein distributions during early adhesion were observed using a confocal microscope (FV3000; Olympus) with a 60× objective.

### 2.4 Data analysis

At *T*_inc_ = 2 and 5 min, 11 and 9 cells, respectively, were examined from one dish in one independent experiment at each time point. All cells at these time points were classified as spherical cells at the earliest stage of adhesion (Fig. S2a,b), and no paxillin localization specific to the adhesion plane was observed. These cells were therefore used only to measure the experimental cell radius. At *T*_inc_ = 15 min, 48 cells from four dishes obtained in two independent experiments were initially examined. Twelve cells showing advanced spreading morphology, including marked flattening, pronounced lamellipodia, or multiple prominent filopodia, were excluded (Figs. S2c, and S3). The remaining 36 cells (7–11 cells per dish) were included in both the binarized paxillin-positive-region analysis and the radial intensity-profile analysis (Fig. 2).

**Figure 2:**
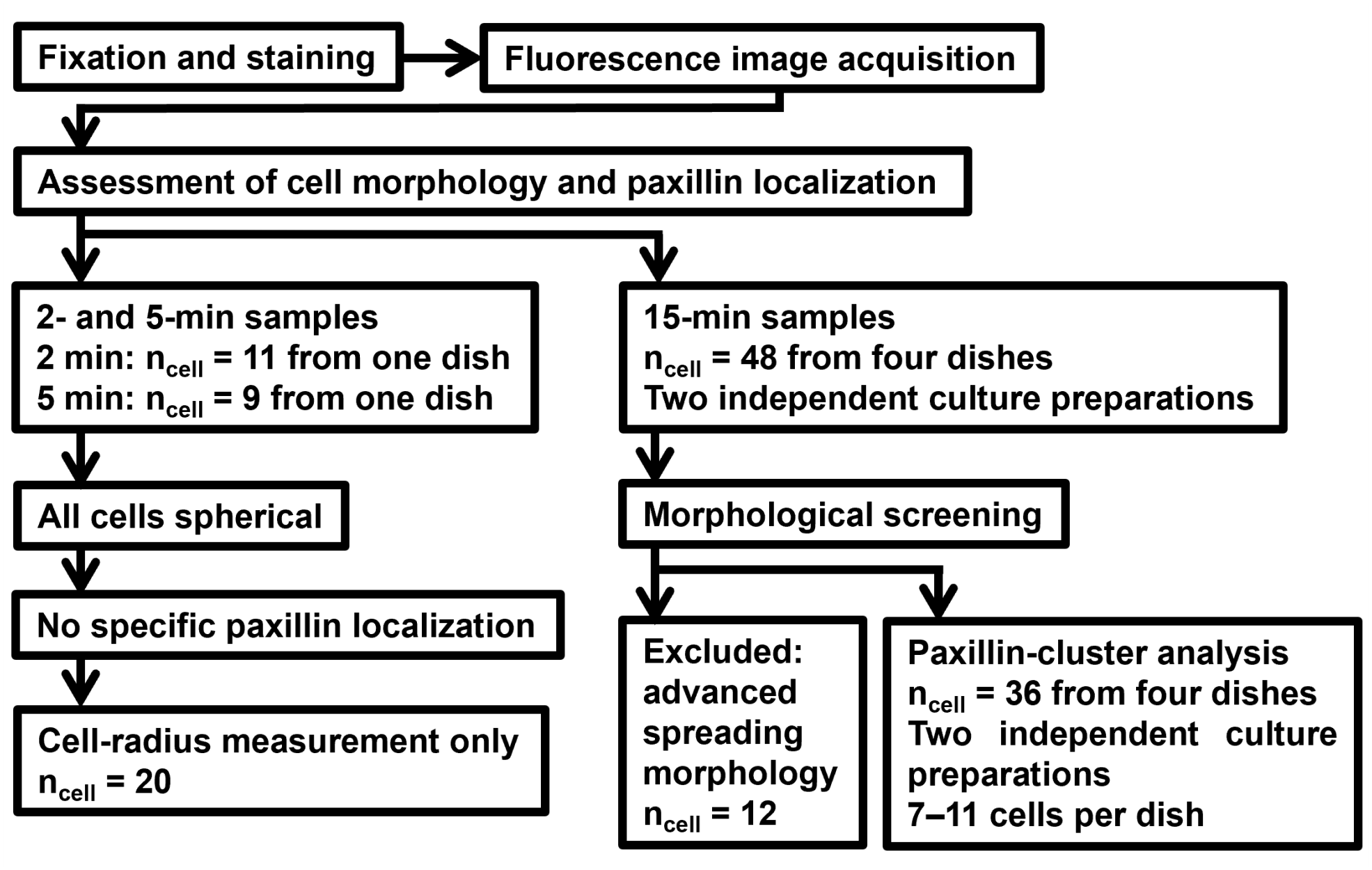
Experimental workflow and sample selection. At 2 and 5 min, 11 and 9 cells, respectively, from one dish and one independent experiment at each time point were used for cell-radius measurement. At 15 min, 48 cells from four dishes obtained in two independent experiments were screened, and 36 cells were included in the paxillin analysis after morphological selection.

Simulated bound integrins and experimentally observed paxillin-positive structures represent distinct molecular observables and cannot be regarded as biologically equivalent. Nevertheless, previous studies support their spatial and functional association during early adhesion (Laukaitis et al., 2001; Wiseman et al., 2004; Yu et al., 2011; Theodosiou et al., 2016). Accordingly, paxillin fluorescence was used here as a spatial readout of early integrin-based adhesion sites.

Experimental paxillin fluorescence images and simulated bound-integrin distributions were analyzed using two complementary approaches. Radial intensity profiles were calculated from the fluorescence images to compare their spatial intensity patterns, whereas cluster metrics were quantified from the corresponding binarized images.

For the simulation results, pseudo-fluorescence images were generated from the coordinates of bound integrins using a point spread function represented by a two-dimensional Gaussian distribution (Fig. 3a). The intensity distribution *I*(*x, y*) used to generate the pseudo-fluorescence images is given by

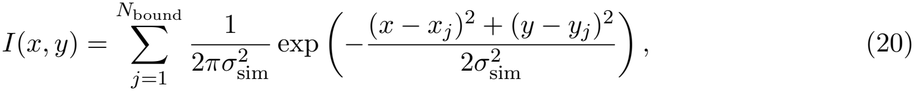

where *N*_bound_ is the number of bound integrins, *x*_*j*_ and *y*_*j*_ represent the position of the *j*-th bound integrin, and *σ*_sim_ is the standard deviation of the two-dimensional Gaussian function. The standard deviation of the Gaussian function, which determines the extent of intensity spreading, was set as follows. The experimental value *σ*_exp_ (∼ 80.84 nm) was estimated using the objective lens with a numerical aperture of 1.42 and the imaging wavelength (∼ 530 nm). The experimental cell radius was measured from spherical cells observed at *T*_inc_ = 2 min and 5 min, and the average value was adopted 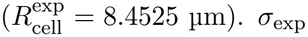 was then scaled by the cell-radius ratio between the simulation and the experiments, 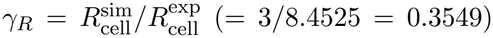, resulting in *γ*_*R*_*σ*_exp_ = 28.69 nm. Based on this estimate, *σ*_sim_ = 28 nm was adopted for the point spread function used to analyze the simulation data. The sensitivity of the cluster-analysis results to *σ*_sim_ was examined by repeating the analysis with *σ*_sim_ = 26 and 30 nm (Fig. S4).

**Figure 3:**
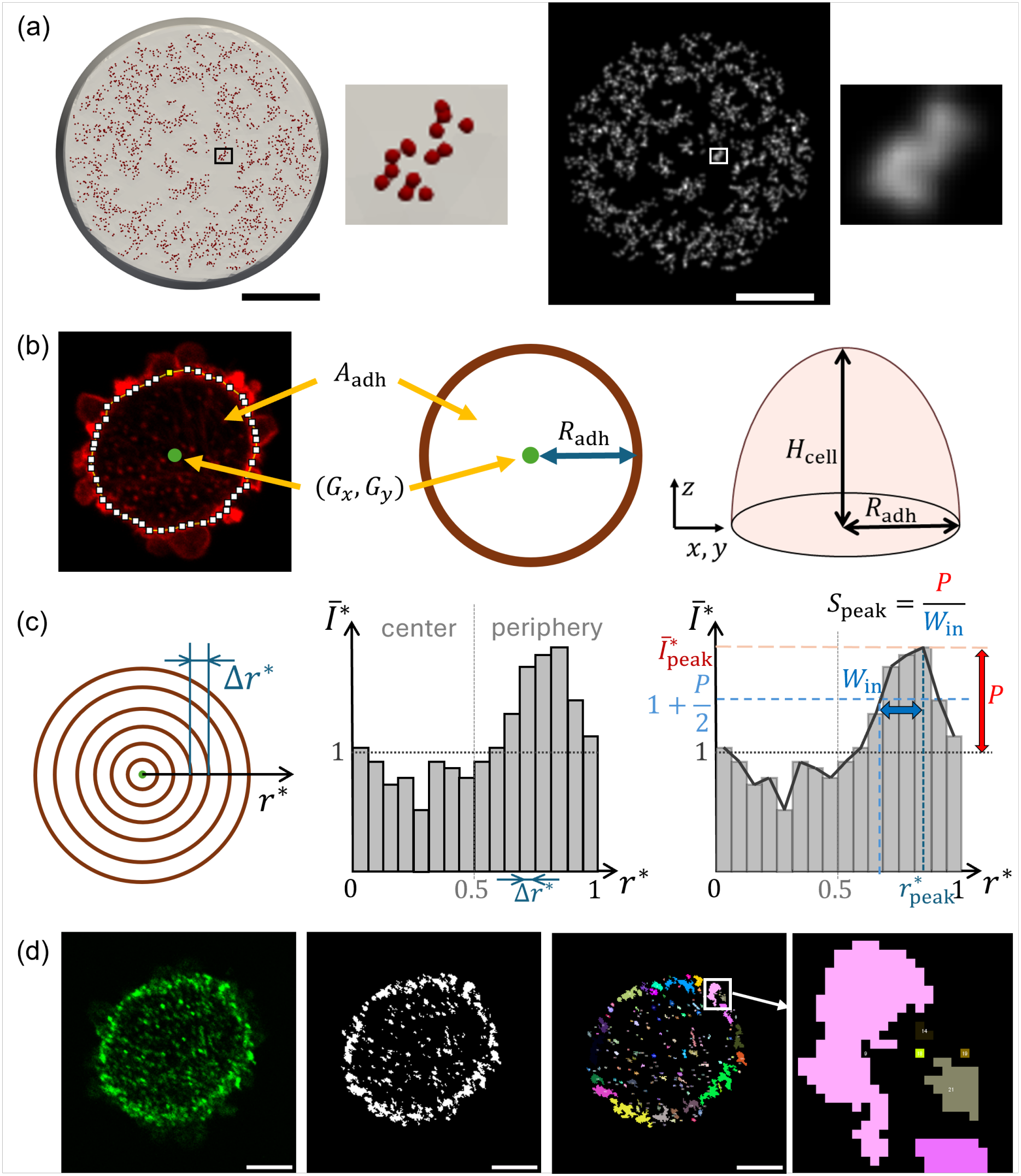
Data analysis methods. **a** The coordinates of bound integrins obtained from the simulation (left) were converted into a pseudo-fluorescence image (right) using a point spread function represented by a two-dimensional Gaussian distribution (*σ*_sim_ = 28 nm). **b** Actin fluorescence images acquired at the cell–substrate interface were used to measure the adhesion area and to calculate the radius of an equivalent-area circle, defined as the adhesion radius *R*_adh_, and the centroid of the adhesion region (left). Early adherent cells in the experiments were approximated as hemi-spheroids, and their surface areas were calculated from the adhesion radius and cell height (right). **c** Concentric annular regions with a normalized radial width of Δ*r*^∗^ = 0.05 were defined within the equivalent-area circle of the adhesion region (left) to evaluate the radial intensity profile. The normalized mean intensity was calculated for each annular interval (center), and the adhesion region was divided into central and peripheral regions at *r*^∗^ = 0.5. The sharpness of the peripheral intensity peak was calculated from the peak height and inward half-width (right). **d** Workflow for analyzing paxillin-positive regions. The paxillin fluorescence image (left) was binarized using the Moments thresholding method (center), and the number and area of each paxillin-positive region were measured (right). Scale bars indicate a normalized length of 0.5.

To establish a common spatial scale for comparing the experimental fluorescence images and simulated pseudo-fluorescence images, a cell-specific reference length *L*_*i*_ was introduced. The spatial scale of each image was normalized by *L*_*i*_, which was defined as

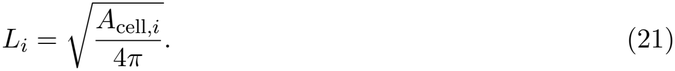

The cell surface area *A*_cell,*i*_ was also used to normalize the metrics in the cluster analysis described below. For the simulation, 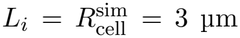. For the experimental images, the basal optical section at which the paxillin fluorescence intensity was maximal along the *z*-axis profile was defined as the cell–substrate adhesion plane. The cell-body adhesion region was manually delineated in Fiji (Schindelin et al., 2012) from the actin fluorescence image at this plane (Fig. 3b). When the boundary was difficult to determine, orthogonal views and the corresponding paxillin fluorescence distribution were also examined. Actin-positive regions judged from these images to originate from bleb-like membrane protrusions were excluded from the cell-body adhesion region. The adhesion area *A*_adh,*i*_ and centroid were measured from the delineated region, and the adhesion radius *R*_adh,*i*_ was defined as the radius of a circle with the same area. For the experimental samples, the cell surface area *A*_cell,*i*_ was estimated by approximating each cell as a hemispheroid using the cell height *H*_cell,*i*_ and adhesion area *A*_adh,*i*_ obtained from the actin fluorescence images:

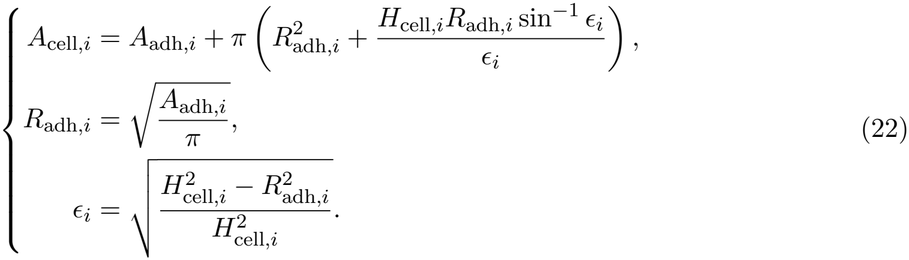

To quantitatively compare the spatial distributions of fluorescence intensity, radial intensity profiles were analyzed using the experimental paxillin fluorescence images and the simulated integrin pseudo-fluorescence images before binarization (Fig. 3c). The centroid of the adhesion region was taken as the origin, and the mean intensity was calculated within concentric annular regions with a normalized radial width of Δ*r*^∗^ = 0.05 for both the experimental and simulation data. The radial distance *r* from the origin was normalized by the adhesion radius *R*_adh,*i*_ as *r*^∗^ = *r/R*_adh,*i*_. The normalized mean intensity in the *k*-th annular region was defined as

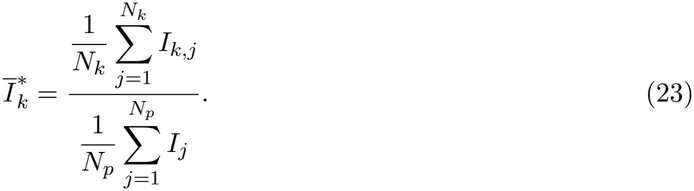

where *N*_*k*_ is the number of pixels in the *k*-th annular region, *I*_*k,j*_ is the intensity of its *j*-th pixel, *N*_*p*_ is the total number of pixels within the adhesion region (0 ≤ *r*^∗^ *<* 1), and *I*_*j*_ is the intensity of its *j*-th pixel.

The fluorescence intensity in the peripheral region relative to that in the central region was quantified using *Q*_periphery_, defined as the ratio of the mean intensity in the peripheral region (0.5 ≤ *r*^∗^ *<* 1) to that in the central region (0 ≤ *r*^∗^ *<* 0.5):

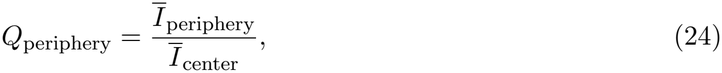

where Ī_center_ and Ī_periphery_ are the mean intensities of all pixels in the central and peripheral regions, respectively. A value of *Q*_periphery_ *>* 1 indicates a peripheral bias, with higher mean intensity in the peripheral region than in the central region.

The sharpness of the intensity peak in the peripheral region was quantified using *S*_peak_. The maximum normalized mean intensity 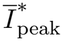 and its radial position 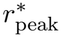 were determined within 0.5 ≤ *r*^∗^ *<* 1. The peak height *P*, inward half-width *W*_in_, and peak sharpness *S*_peak_ were defined as

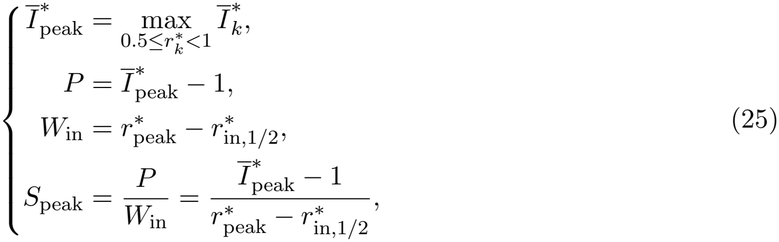

where 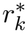 is the normalized radial position of the *k*-th annular region and 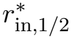 is the radial position on the inward side of the peak at which the normalized mean intensity reaches 1 + *P/*2. A larger *S*_peak_ indicates a more sharply rising peripheral intensity peak.

For the cluster-based comparison, the experimental fluorescence and simulated pseudo-fluorescence images were binarized using the Moments thresholding method to extract paxillin-positive and bound-integrin-rich regions, respectively (Fig. 3d). The resulting binarized images were analyzed using the Analyze Particles function in Fiji to quantify the number and area of the extracted regions. Hereafter, the regions extracted by binarization from both datasets are referred to simply as clusters. Using the cell surface area *A*_cell,*i*_ defined above, the following normalized cluster metrics were calculated: the normalized adhesion area 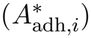, cluster number density 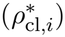, total cluster area fraction within the adhesion region 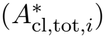, and normalized mean cluster area 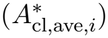:

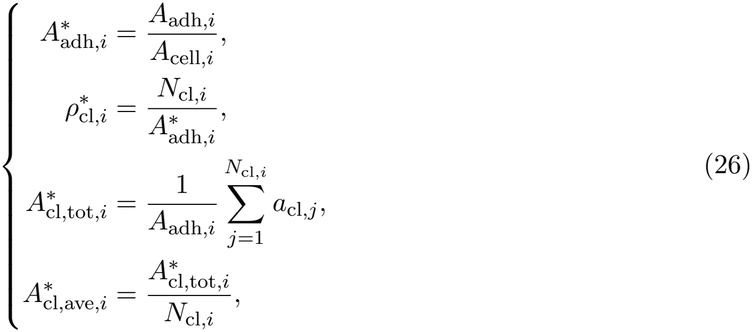

where *N*_cl,*i*_ is the number of clusters within the adhesion region, and *a*_cl,*j*_ is the area of the *j*-th cluster.

## 3 Results

### 3.1 Time evolution of simulated cell adhesion

To provide an overview of the simulated adhesion process, representative results are shown for the reference condition (*λ* = 1) and a lower-stiffness condition (*λ* = 0.1). In the latter condition, 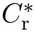 was also set to 0.1 times its reference value to approximately preserve the mechanical relaxation timescale. The cell membrane adhered to the substrate surface, and the adhesive area increased over time (Fig. 4a, b). Compared with the reference condition (*λ* = 1), the cell spread more extensively and adopted a different shape at *λ* = 0.1. Integrin binding did not occur immediately upon initial cell membrane–substrate contact but began shortly thereafter, with the number of bound integrins subsequently increasing over time. In the early stage, single integrin binding events were frequently observed in the periphery of the adhesion area. Moreover, additional integrins clustered around already-bound integrins, and the number of these clusters increased over time. Integrin clusters were more frequently observed in the periphery than in the center of the adhesion area. By *t*^∗^ = 0.417, the normalized adhesion area had approached a plateau. Nevertheless, integrins continued to cluster around pre-existing bonds, and the bound fraction *N*_bound_*/N*_I_, where *N*_bound_ is the number of bound integrins, continued to increase (Fig. 4c). The normalized time at which these two quantities coincided was defined as 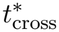, and the integrin cluster distribution and the distance between the membrane and substrate surfaces were evaluated at 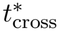 (Fig. 4d–f). As discussed earlier, integrin clusters tended to be more abundant in the periphery than in the center of the adhesion area (Fig. 4d). Visualization of the distance between the membrane and the substrate surface revealed regions where the membrane locally approached the substrate (Fig. 4e), and these regions corresponded to the distribution of integrin clusters (Fig. 4f).

**Figure 4:**
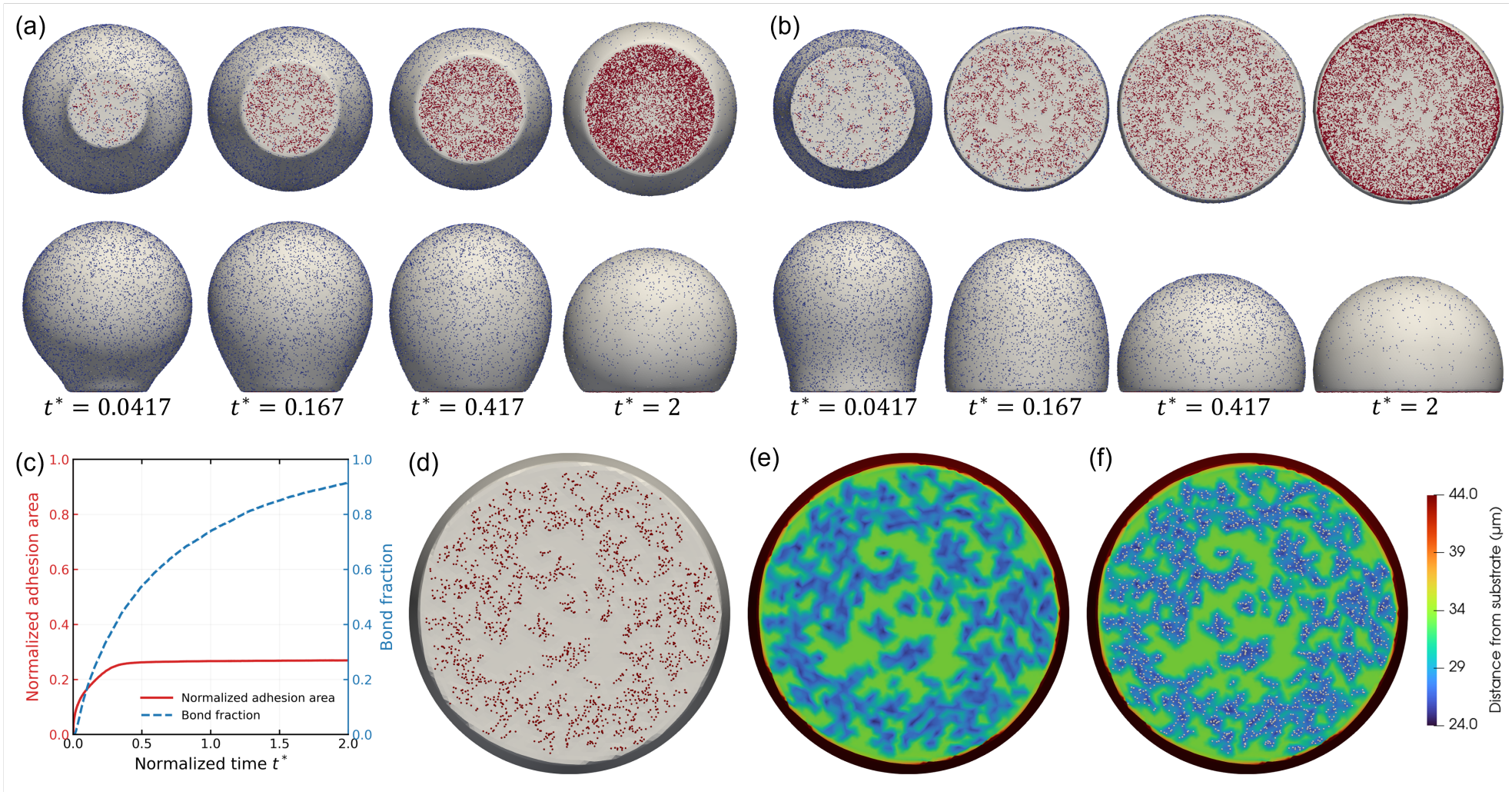
Results of a cell adhesion simulation. Time evolution of the adhesive cell and integrins for *λ* = 1 (**a**) and *λ* = 0.1 (**b**). In each panel, the top and bottom rows show the cell and integrins as observed from below and from the side, respectively. Panels **c**–**f** show the results for *λ* = 0.1. Integrins distributed on the membrane are shown as particles, and inactive, active but unbound, and bound integrins are shown in blue, white, and red, respectively. **c** Time evolution of the normalized adhesion area and bound fraction. **d**–**f** The bottom of the cell at 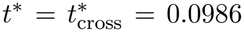 when the normalized adhesion area and bound fraction coincide. **d** Bound integrins are visualized only as red points. **e** Only the membrane is shown colored by distance from the substrate. **f** Merged image of **d** and **e**, where white points show bound integrins.

### 3.2 Experimental observations and radial distribution of paxillin

Figure 5 shows the experimental results. In cells at the early stage of adhesion, blebs (balloon-like membrane protrusions) were observed near the adhesion plane, whereas thick fibrous structures were not detected (Fig. 5a). Paxillin was localized at the adhesion plane and, in general, more paxillin clusters were observed in the periphery of the adhesion region. Meanwhile, some samples exhibited band-like paxillin clusters along the edge of the adhesion region (Fig. 5b, left), whereas in others paxillin clusters were sparsely distributed near the edge of the adhesion region (Fig. 5b, right), indicating variability in the distribution patterns of paxillin clusters in the peripheral region. In addition, some samples exhibited paxillin clusters in the center of the adhesion area at a density comparable to that in its periphery (Fig. 5b, center). These results indicate that the distribution of paxillin clusters varied substantially among samples, even when the normalized adhesion areas were similar.

**Figure 5:**
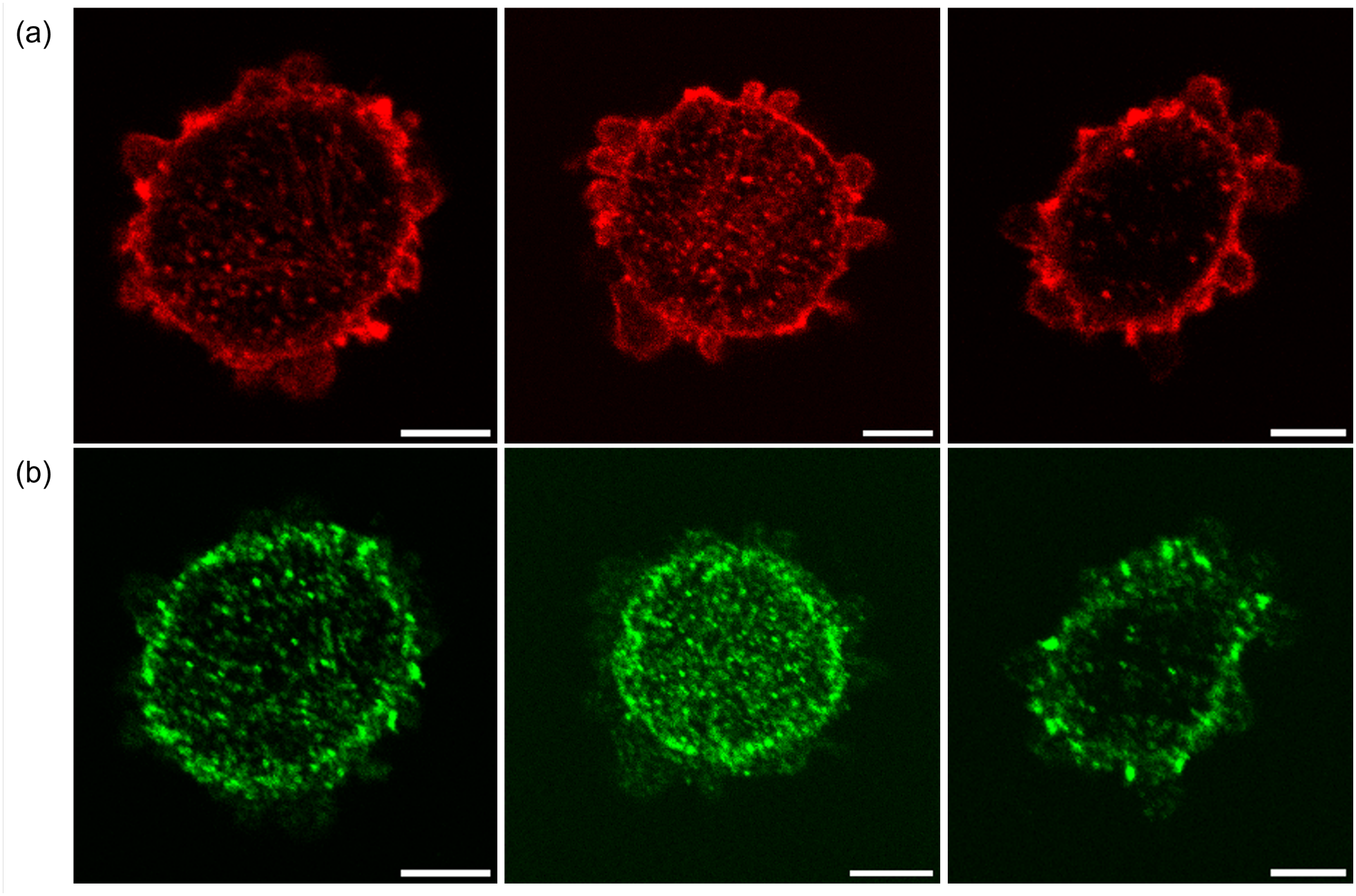
Results of cell adhesion observation. Representative images were selected from the analyzed dataset of 36 cells from four dishes obtained in two independent experiments. Fluorescence images of actin (**a**) and paxillin (**b**). Different clustering distributions appear even at similar normalized adhesion areas, 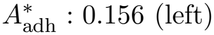, 0.132 (center), and 0.124 (right). Scale bars indicate a normalized length of 0.5.

To examine this sample-to-sample variability across all 36 cells included in the analysis, we evaluated the spatial distribution of paxillin. The paxillin fluorescence images were divided into six groups (sextiles) based on the normalized adhesion area 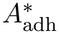 and, within each group, arranged in ascending order of *Q*_periphery_ (Fig. 6a, and Fig. S5). Specifically, the image in the *k*-th row and *l*-th column represents a cell belonging to the *k*-th sextile of 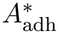 and having the *l*-th smallest *Q*_periphery_ within that sextile. The corresponding radial profiles of normalized mean intensity are shown in the same arrangement in Fig. 6b and Fig. S6. Most cells exhibited a peripheral bias, with higher mean fluorescence intensity in the peripheral region than in the central region. However, even among cells with comparable 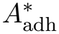, some exhibited paxillin clusters in the central region and a gradual radial change in intensity, whereas others showed strong paxillin localization at the edge of the adhesion region and a sharp peripheral intensity peak. Notably, a subset of cells in the lower 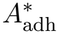 sextiles exhibited higher mean fluorescence intensity in the central region than in the peripheral region.

**Figure 6:**
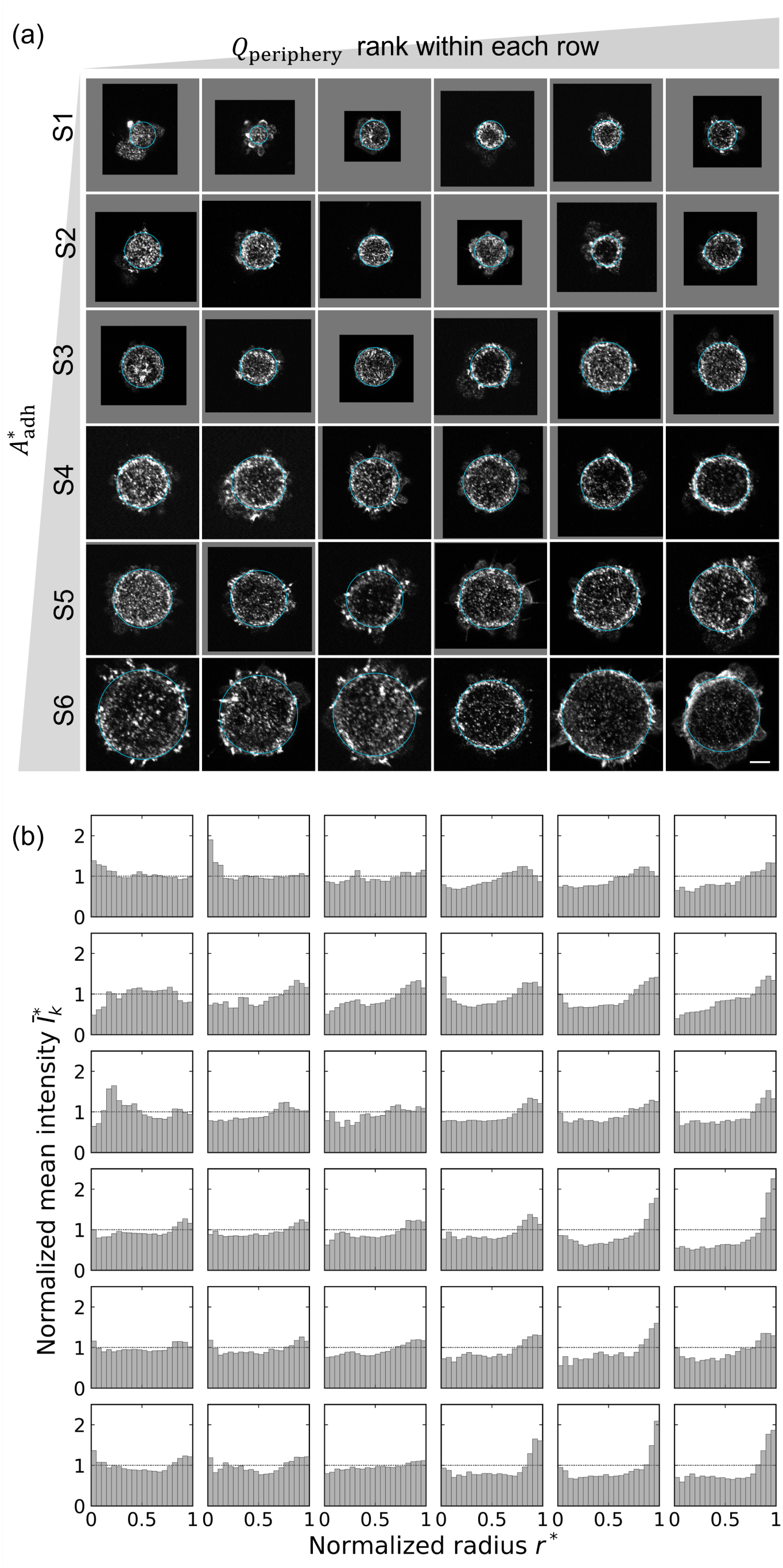
Spatial distribution of paxillin during early adhesion. All 36 analyzed cells from four dishes obtained in two independent experiments are shown. Fluorescence images of paxillin (**a**) and their radial distributions of normalized mean intensity (**b**). In **b**, the distributions are shown as bar plots, with each bar representing the normalized mean intensity within an annular region of normalized radial width Δ*r*^∗^ = 0.05. Blue circles indicate the adhesive radius. The scale bar indicates a normalized length of 0.5.

### 3.3 Effects of membrane stiffness on simulated adhesion and integrin distribution

To examine how membrane stiffness affects adhesion and integrin distribution during early adhesion, the membrane-stiffness-related coefficients were jointly scaled from their reference values using *λ* = 10, 1, 0.2, and 0.1. To keep the mechanical relaxation timescale approximately constant across these conditions, 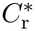 was also set to *λ* times its reference value, 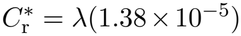, as follows from Eq. (11). Five independent simulations with different random seeds were performed for each condition. For each membrane-stiffness condition, pseudo-fluorescence images were generated from the simulated distributions of bound integrins, and their radial intensity profiles were evaluated using the same procedure as for the experimental images. In Fig. 7a, each row represents a membrane-stiffness condition, and each column represents a successive snapshot. The corresponding radial profiles of normalized mean intensity are shown in the same arrangement in Fig. 7b. At corresponding time points, the normalized adhesion area was larger under lower membrane-stiffness conditions. Although bound-integrin clusters were also observed in the central region during the early stages of adhesion, the relative intensity in the peripheral region increased as adhesion progressed; under lower membrane-stiffness conditions, aggregates formed along the edge of the adhesion region at later stages. Under all membrane-stiffness conditions examined, bound integrins tended to be distributed more densely in the peripheral region than in the central region of the adhesion area.

**Figure 7:**
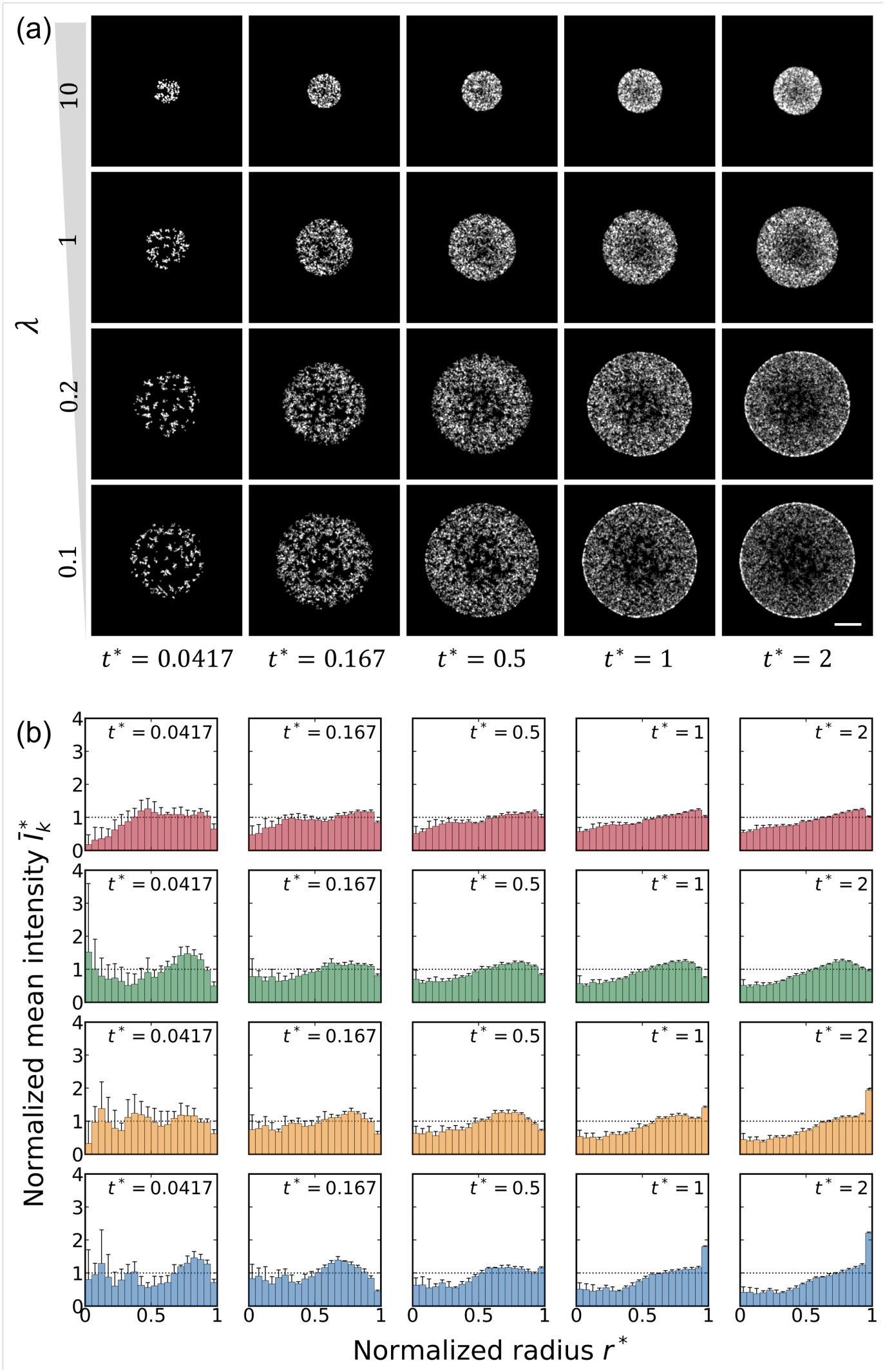
Spatial distribution of bound integrins in the cell adhesion model. Pseudo-fluorescence images of integrin (**a**) and their radial distributions of normalized mean intensity (**b**). In **a** and **b**, the rows correspond to different values of *λ*, and the columns show time evolution. In **b**, the distributions are shown as bar plots with a normalized radial bin width of Δ*r*^∗^ = 0.05. Five simulations with different random seeds were performed for each condition; the bars show the mean across the five simulations, and the upper error bars indicate the corresponding standard deviation across the five random seeds. The scale bar indicates a normalized length of 0.5.

### 3.4 Comparison of spatial distributions and cluster metrics between experiments and simulations

The radial distributions of normalized mean intensity obtained from the experimental fluorescence images and simulated pseudo-fluorescence images were compared using *Q*_periphery_ and *S*_peak_, as defined in Section 2.4 and Eqs. (24) and (25), respectively (Fig. 8, Fig. S7ab, and Fig. S8ab).

**Figure 8:**
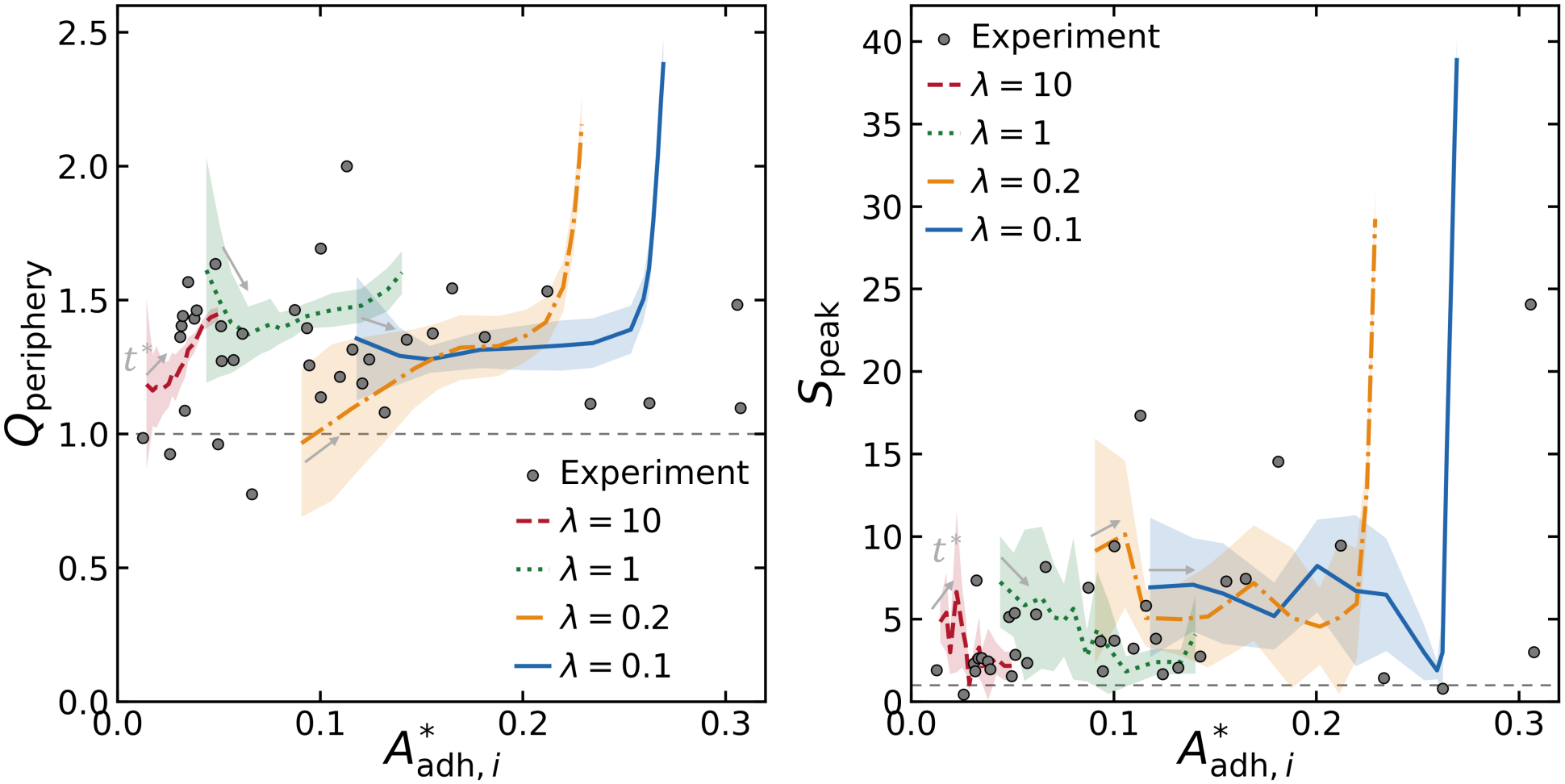
Comparison of intensity distribution between simulation and experimental results. The horizontal axis represents the normalized adhesion area 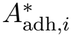 the vertical axes show the ratio of the peripheral to central mean intensity *Q*_periphery_ (left) and the sharpness of the peripheral intensity peak *S*_peak_ (right). Gray circles indicate the experimental data, and the colored lines and shaded bands show the time evolution of the simulation results for different values of *λ*. The experimental data comprise 36 cells from four dishes obtained in two independent experiments. Five simulations with different random seeds were performed for each membrane-stiffness condition. The lines show the mean across the five simulations, and the shaded bands indicate the corresponding mean ± standard deviation across the five random seeds. Gray arrows indicate the direction in which *t*^∗^ increases.

In the experiments, 32 of the 36 cells (88.9%) exhibited a peripheral bias (*Q*_periphery_ *>* 1). Although *Q*_periphery_ varied among cells, it showed no clear change with the normalized adhesion area 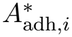 (Fig. 8, left). In contrast, *S*_peak_ showed a slight tendency to increase with 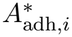, although substantial variability was observed among cells (Fig. 8, right).

Most simulated profiles also exhibited a peripheral bias (*Q*_periphery_ *>* 1; 269 of 280 profiles). The simulated *S*_peak_ values followed condition-dependent time courses and were within the experimental range at many time points. In the five-seed mean trajectories under the higher membrane-stiffness conditions (*λ* = 10 and 1), *Q*_periphery_ generally increased with 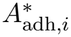, whereas *S*_peak_ tended to decrease and subsequently fluctuate. Consistent with these trends, as the adhesion region expanded, the peripheral bias generally became stronger, while the radial intensity profiles showed a relatively gradual increase toward the periphery. During this progression, both metrics remained within the range of the experimental data. Under the lower membrane-stiffness conditions (*λ* = 0.2 and 0.1), by contrast, *Q*_periphery_ and *S*_peak_ continued to increase after 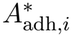 had nearly plateaued. This late-stage increase indicates that the accumulation of bound integrins along the edge of the adhesion region became stronger and that the peripheral intensity peak became more pronounced and sharper, consistent with the temporal evolution shown in Fig. 7.

Cluster metrics obtained from the experimental paxillin fluorescence images and the simulated pseudo-fluorescence images were compared (Fig. 9, Fig. S7c–e, and Fig. S8c–e).

**Figure 9:**
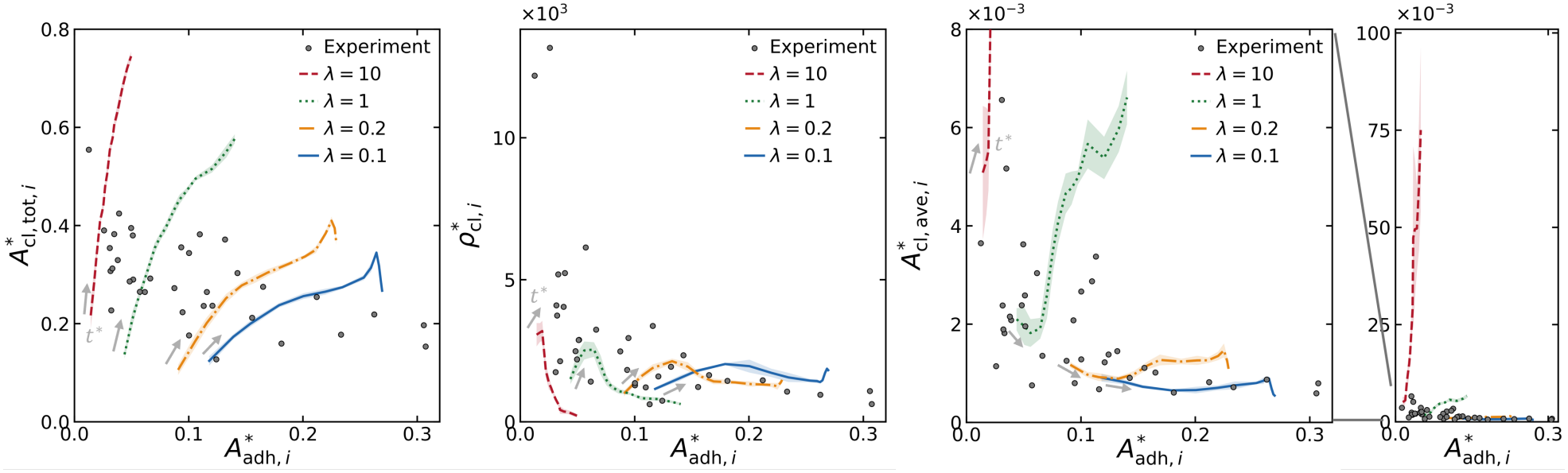
Comparison of clusters between simulation and experimental results. The horizontal axis represents the normalized adhesion area 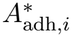 the vertical axes show the total cluster area fraction ^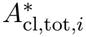^, normalized cluster number density 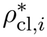 (center), and normalized mean cluster area 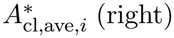. Gray circles indicate the experimental data, and the colored lines and shaded bands show the time evolution of the simulation results for different values of *λ*. The experimental data comprise 36 cells from four dishes obtained in two independent experiments. Five simulations with different random seeds were performed for each membrane-stiffness condition. The lines show the mean across the five simulations, and the shaded bands indicate the corresponding mean ± standard deviation across the five random seeds. Gray arrows indicate the direction in which *t*^∗^ increases.

In the experiments, the total cluster area fraction 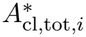, cluster number density 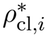, and normalized mean cluster area 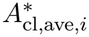 all tended to decrease as 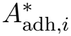 increased. The total cluster area fraction decreased gradually while retaining substantial cell-to-cell variability. By contrast, the cluster number density and normalized mean cluster area varied widely at small 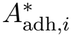, whereas both their values and variability decreased as 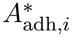 increased.

In the simulations, the total cluster area fraction initially increased as 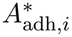 increased. Because expansion of the adhesion region preceded cluster formation to a greater extent under lower membrane-stiffness conditions, the total cluster area fraction was smaller at lower membrane stiffness when compared at the same 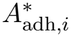.

For *λ* = 0.1, the total cluster area fraction decreased at the final stage of the time series. This decrease may reflect, at least in part, the effects of pseudo-fluorescence image generation and binarization rather than a direct physical decrease in the area occupied by clustered integrins. As integrins accumulated along the edge of the adhesion region, the local density of bound integrins increased, producing high-intensity regions near the edge after application of the point spread function. Consequently, the intensity distribution and binarization threshold changed, which may have excluded relatively low-intensity regions from the detected clusters and reduced the total cluster area fraction.

The total cluster area fraction obtained from the simulations was of the same order of magnitude as that in the experiments (Fig. 9, left). Furthermore, under the conditions *λ* ≤ 1, both the cluster number density and normalized mean cluster area decreased as the normalized adhesion area increased, consistent with the trends observed in the experiments (Fig. 9, center and right).

### 3.5 Effect of integrin–ligand bond strength on membrane deformation and integrin clustering

Figure 10 shows the simulation results at 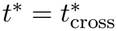 obtained by varying the dimensionless coefficient of the integrin–ligand bond force as 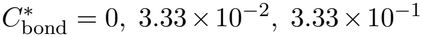, and 3.33, corresponding to 0, 0.1, 1, and 10 times its reference value, respectively. In all conditions, the membrane-stiffness-related coefficients were set to *λ* = 0.1, and 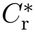 was set to 0.1 times its reference value (1.38 × 10^−6^). Five independent simulations with different random seeds were performed for each condition. Without the integrin–ligand bond force 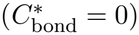, distinct localized high-density integrin clusters scarcely formed. A peripheral bias was observed under all bond-force conditions, although its strength varied (Fig. 10c). The number density of bound integrins in each binarized cluster was calculated as *N*_int,*c*_*/A*_cl,*c*_, where *N*_int,*c*_ and *A*_cl,*c*_ are the number of bound integrins and the area of cluster *c*, respectively. The area-weighted mean across the detected clusters was then calculated as ⟨*N*_int,*c*_*/A*_cl,*c*_⟩_*A*_. Among the conditions with a nonzero bond force, a smaller bond-force contribution produced more localized membrane deformation and cluster formation and resulted in a higher ⟨*N*_int,*c*_*/A*_cl,*c*_⟩_*A*_ (Fig. 10d).

**Figure 10:**
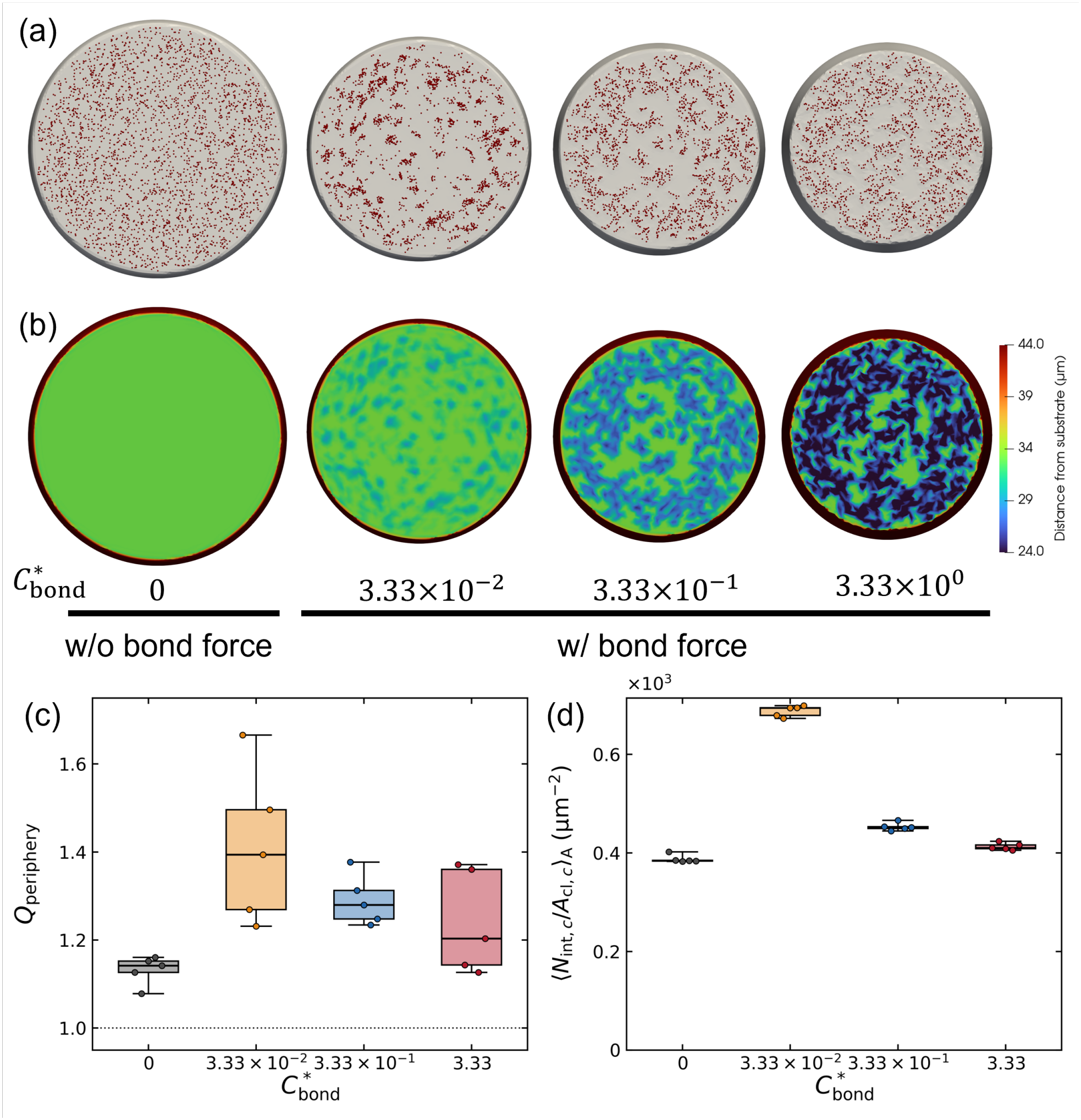
Effect of the integrin–ligand bond force on integrin clustering. **a** Bottom views of the adhered membrane at 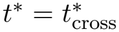, when the normalized adhesion area and bound fraction coincide. From left to right, 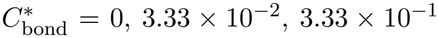, and 3.33; red dots represent bound integrins. The corresponding values of 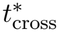 are 0.3691, 0.1555, 0.09857, and 0.08300, respectively. **b** Membrane–substrate distance. **c, d** Box plots of *Q*_periphery_ and the area-weighted number density of bound integrins within the detected clusters ⟨*N*_int,*c*_*/A*_cl,*c*_⟩_*A*_, respectively, at 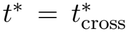. Five simulations with different random seeds were analyzed.

### 3.6 Effect of spreading speed on integrin clustering

Figure 11 shows the simulation results at 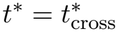 obtained by varying the dimensionless damping coefficient as 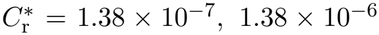, and 1.38 × 10^−5^, corresponding to 0.01, 0.1, and 1 times its reference value, respectively, to investigate the effect of adhesion-region spreading speed on integrin clustering. Throughout this analysis, the membrane-stiffness-related coefficients were fixed at *λ* = 0.1. Five independent simulations with different random seeds were performed for each condition. Under the fast-spreading condition, numerous integrin binding events occurred over a wide area of the adhesion region, and many small clusters were sparsely distributed at 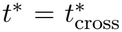 (Fig. 11b, e). The corresponding radial distribution showed *Q*_periphery_ *>* 1, indicating that bound integrins were biased toward the periphery (Fig. 11d). Under the slow-spreading condition, a small number of clusters initially formed preferentially in the periphery (Fig. 11c) and developed into larger clusters by 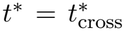 (Fig. 11e). At this time point, *Q*_periphery_ was lower than under the fast-spreading condition, indicating a relatively weaker peripheral bias (Fig. 11d).

**Figure 11:**
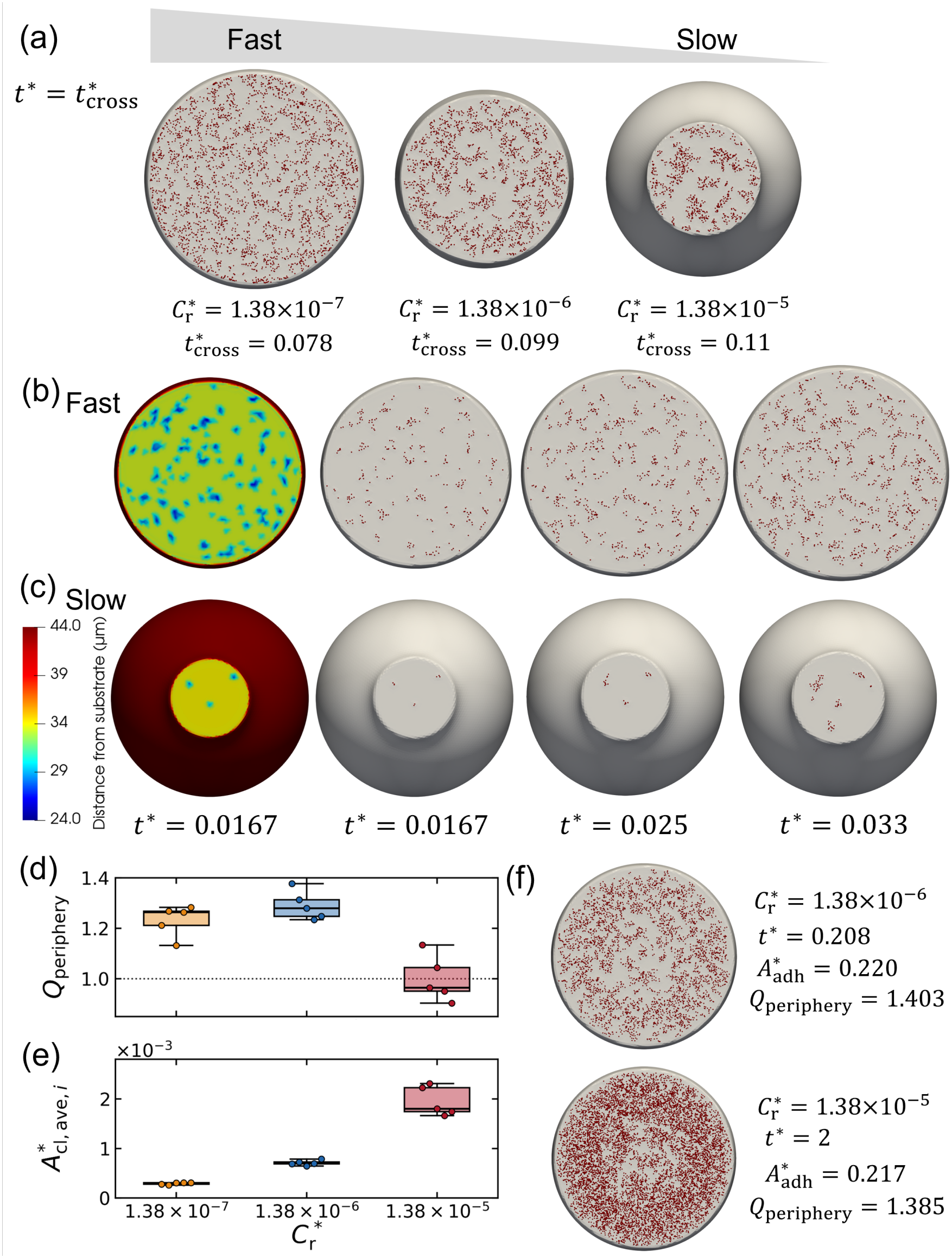
Effect of the spreading speed on integrin clustering. **a** Bottom views of the adhered membrane at 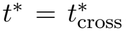, when the normalized adhesion area and bound fraction coincide. From left to right, 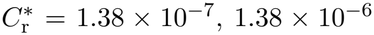, and 1.38 × 10^−5^ correspond to decreasing spreading speed; red dots represent bound integrins. The corresponding values of 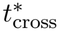 are 0.0784, 0.0986, and 0.1100, respectively. **b, c** Early dynamics of integrin binding under the fast- and slow-spreading conditions, respectively. The leftmost panel in each row shows the membrane–substrate distance. **d, e** Box plots of *Q*_periphery_ and the normalized mean cluster area 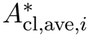, respectively, at 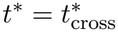. Five simulations with different random seeds were analyzed. **f** Distributions of bound integrins at nearly identical 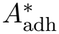 and 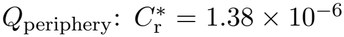 at *t*^∗^ = 0.2083 (top) and 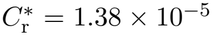 at *t* = 2.0 (bottom). Scale bars indicate a normalized length of 0.5.

We further compared later states with nearly identical 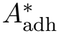 and 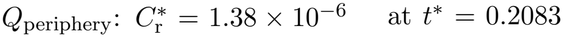 and 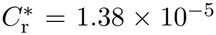at *t*^∗^ = 2.0. In the former, discrete clusters were scattered around the periphery, whereas in the latter, a band-like distribution had formed along the periphery (Fig. 11f).

## 4 Discussion

### 4.1 Comparison with experimental observations

The present model was developed to clarify how cellular-scale integrin clustering emerges by coupling membrane mechanics with mesoscopic integrin dynamics. The simulations showed that integrin– ligand binding induced local membrane deformation and promoted subsequent binding, thereby forming clusters of bound integrins. Examination of the bound-integrin particle distributions showed that clusters formed at high density in the periphery as the adhesion region expanded and, under the low-stiffness conditions, further developed into aggregated structures along the edge of the adhesion region over time (Fig. 7). This tendency was also demonstrated quantitatively by evaluating the radial intensity distributions of pseudo-fluorescence images generated from the particle distributions (Fig. 8). A similar peripheral bias was observed in the experimental paxillin distributions (Figs. 6 and 8).

The experimental paxillin fluorescence images showed different spatial distributions even among cells with comparable normalized adhesion areas 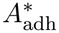, whereas cells with different 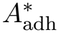 sometimes showed similar spatial distributions (Fig. 6a). In most cells, *Q*_periphery_ *>* 1, indicating that paxillin fluorescence was biased toward the peripheral rather than the central region of the adhesion area (Fig. 8, left). Qualitative inspection of the images suggested that accumulation in the periphery or along the edge of the adhesion region tended to be more evident in cells with relatively large 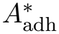. However, *Q*_periphery_ showed no clear change with 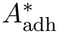, whereas *S*_peak_ showed a slight tendency to increase, with substantial variability among cells (Fig. 8). Overall, the cluster metrics tended to decrease with increasing 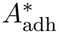, although the total cluster area fraction retained substantial cell-to-cell variability (Fig. 9, left). The cluster number density and normalized mean cluster area varied widely, particularly among cells with small 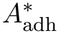 (Fig. 9, center and right). All cells analyzed in this study were fixed and stained 15 min after seeding; however, the interval between the initial contact of each cell with the substrate and fixation was not necessarily identical among cells. Thus, cell-to-cell differences in the onset of adhesion may have contributed to the variability in 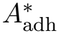 and to the formation of different paxillin distributions at comparable 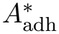. However, because 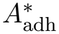 also depends on the mechanical state of the membrane and morphological variability among cells, it does not correspond one-to-one with the elapsed time after adhesion initiation.

In the simulations, *Q*_periphery_ *>* 1 was also observed at most time points, reproducing the peripheral bias observed in the experimental paxillin fluorescence. Moreover, under each membrane-stiffness condition, *Q*_periphery_ and *S*_peak_ were within ranges comparable to the experimental values at many time points (Fig. 8). Under the high-stiffness conditions, *S*_peak_ decreased after the initial transient period, resulting in a relatively gradual peripheral distribution. Under the low-stiffness conditions, by contrast, both *Q*_periphery_ and *S*_peak_ continued to increase after 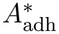 had nearly reached a steady value, indicating further accumulation along the edge of the adhesion region (Figs. 7 and 8). This partial agreement with the experiments was also observed in the cluster metrics. For each cluster metric, the simulation results under the individual membrane-stiffness conditions fell within the overall range of the experimental data at some time points (Fig. 9). Under the lower-stiffness conditions, the decreases in cluster number density and normalized mean cluster area with increasing 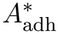 were also similar to the experimental trends. At later time points, however, some of the metrics moved outside the experimental range. Thus, the simulations under the individual membrane-stiffness conditions reproduced parts of the ranges and trends observed experimentally, but did not reproduce the full cell-to-cell variability.

One factor that may have contributed to this discrepancy is that membrane stiffness was held temporally constant in the simulations. Early cell spreading has been suggested to involve bleb formation, a reduction in membrane tension, and the recruitment of membrane reservoirs (Pietuch and Janshoff, 2013). Bleb formation requires local detachment of the membrane from the underlying actomyosin cortex, which provides mechanical support to the membrane (Welf et al., 2020). Because bleb formation was also observed in our experiments (Fig. 5), the mechanical coupling between the membrane and cortex may be locally reduced during early adhesion, accompanied by spatiotemporal variation in the apparent membrane stiffness. However, membrane stiffness was not directly measured in this study and was held temporally constant in each simulation. Therefore, if the apparent membrane stiffness of real cells changes as adhesion progresses, the adhesion process may exhibit behavior that interpolates among the fixed membrane-stiffness conditions examined in this study (Figs. 7–9). Taken together, the cell-to-cell variability observed experimentally may reflect differences in the onset of adhesion as well as cell-to-cell differences and temporal changes in the mechanical state of the membrane.

### 4.2 Mechanism of integrin clustering

Varying membrane stiffness altered both the cell-scale mechanical balance between membrane stiffness and physical adsorption, which determines the adhesion shape and area, and the local mechanical response of the membrane to the bond force. Therefore, comparison among the membrane-stiffness conditions alone could not isolate the mechanism responsible for the formation of individual clusters. To separate these effects, we fixed the parameters governing membrane stiffness and physical adsorption, varied 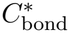, and examined the role of bond-force-induced membrane deformation in cluster formation (Fig. 10).

When integrins bind to ligands, the resulting bond force acts on the membrane and locally brings the membrane closer to the substrate surface near the bound integrins. This locally increases the probability of subsequent binding events. Such a mechanism for the formation of a single integrin cluster has already been demonstrated in a previous computational study using a periodic adhesion surface (Paszek et al., 2009), and is consistent with the results of the present study. Among the conditions with a finite bond-force contribution, a smaller contribution caused membrane deformation to remain more localized around individual bound integrins, thereby restricting the region in which the binding probability was enhanced. At 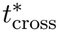, the mean density of bound integrins within clusters was highest at 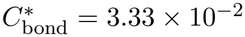 and decreased with increasing 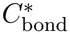 (Fig. 10b, d). By contrast, a larger bond-force contribution extended membrane deformation and the region of enhanced binding over a wider area, resulting in larger detected clusters with a lower number density of bound integrins.

When the bond force did not act on the membrane, the local positive feedback whereby deformation around an existing bond promotes subsequent binding did not operate. Consequently, integrins bound relatively sparsely throughout the adhesion region, and no clearly localized highdensity clusters comparable to those in the other conditions were observed (Fig. 10a). These results indicate that binding-induced membrane deformation promotes the formation of high-density integrin clusters when it has a sufficient magnitude while remaining localized near pre-existing bonds. Previous studies have suggested that NA formation requires coupling to the actin cytoskeleton (Changede et al., 2015). Although the present model does not explicitly include the actin cytoskeleton, the membrane-stiffness parameters may be interpreted phenomenologically as representing the effective mechanical resistance of the membrane–cortex system. Because the membrane-stiffness parameters were fixed in this analysis, varying 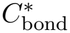 changed the balance between the bond force and the fixed mechanical resistance of the membrane. Therefore, the present results suggest that actin– membrane coupling may influence integrin clustering by altering the effective mechanical properties of the membrane–cortex system and, consequently, the binding-induced membrane deformation.

### 4.3 Mechanism governing the spatial distribution of integrin clusters

At 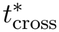, *Q*_periphery_ *>* 1 under all conditions examined by varying 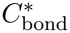, including the condition in which the bond force did not act on the membrane (Fig. 10c). Thus, the initial bias of bound integrins toward the adhesion periphery cannot be explained solely by the local positive feedback arising from binding-induced membrane deformation. During the early stage of adhesion, the membrane– substrate distance was largely flat within the basal adhesion region, although a slight local reduction was observed at the edge of the adhesion region (Fig. 10b). This fine-scale proximity at the edge may have favored initial binding there. Subsequently, the flat basal membrane moved closer to the substrate as a whole, thereby reducing the spatial difference in binding probability within the adhesion region. In addition, integrins in the present model are distributed on the lateral membrane and diffuse over the membrane surface. After the adhesive morphology has developed, integrins moving from the lateral membrane toward the basal adhesion surface may first enter and be captured at the edge of the adhesion region, which separates these membrane regions. Together, the fine-scale membrane–substrate geometry at the edge of the adhesion region and the subsequent supply of integrins from the lateral membrane may establish a baseline peripheral bias even in the absence of bond-force-mediated membrane deformation.

When the bond force acted on the membrane, the peripheral bias tended to be stronger than that in the absence of the bond force (Fig. 10c). Binding events initiated near the edge of the adhesion region locally drew the membrane toward the substrate and thereby promoted subsequent binding in their vicinity, amplifying cluster formation in the peripheral region. Consistent with this interpretation, peripheral bias was observed across the membrane-stiffness conditions, and accumulation along the adhesion edge became progressively more pronounced under the low-stiffness conditions (Figs. 7 and 8). Taken together, these results suggest that the cell-scale spatial distribution of integrin clusters is initially biased by the membrane–substrate geometry and the subsequent supply of integrins from the lateral membrane, whereas binding-induced membrane deformation further promotes cluster formation in the peripheral region.

### 4.4 Effect of time-scale balance by spreading speed

The adhesive shape of the cell membrane is determined by the mechanical balance between membrane stiffness and physical adsorption. However, it remains unclear whether integrin clustering dynamics and the resulting spatial patterns are determined solely by static mechanical equilibrium or are also influenced by spreading speed. In the present model, the spreading speed of the adhesion region is regulated by 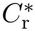, which represents the effective dissipative coupling of the membrane–cortex complex with the intracellular and extracellular environments. We therefore varied 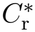 while keeping the membrane-stiffness-related coefficients fixed and examined its effect on integrin clustering dynamics. Under slow-spreading conditions, the overall membrane–substrate distance remained relatively large for a longer period. When an integrin binding event occurred during this period, bond-induced membrane deformation promoted subsequent binding near the existing bond, allowing local cluster formation to proceed. The membrane–substrate distance subsequently decreased and binding proceeded throughout the adhesion region; however, accumulation into the clusters formed earlier continued, resulting in a smaller number of relatively large clusters (Fig. 11c, e). At 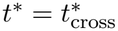, the radial distribution of normalized mean intensity showed a relatively weak peripheral bias (Fig. 11d).

In contrast, under fast-spreading conditions, the membrane–substrate distance became uniformly small before integrin binding had sufficiently progressed (Fig. 11b). Thus, the membrane–substrate distance decreased over a wide area before subsequent binding near existing bonds could progress sufficiently. Integrin binding consequently began at many locations within the adhesion region, producing a larger number of smaller clusters than under the slow-spreading condition (Fig. 11b, e). At 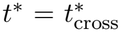, the normalized mean intensity in the peripheral region was higher than that in the central region, indicating a peripheral bias (Fig. 11d).

Moreover, the two later states selected to have nearly identical 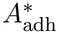 and *Q*_periphery_ exhibited distinct cluster morphologies: discrete clusters were scattered around the periphery under the intermediate-spreading condition, whereas a band-like peripheral distribution formed under the slow-spreading condition (Fig. 11f). This difference suggests that, even at similar normalized adhesion areas and radial biases, the spatial organization of integrin clusters depends on the spreading speed and the preceding formation history.

These results suggest that integrin clustering patterns depend not only on the static mechanical state determined by membrane deformation and physical adsorption, but also on the relative timescales of adhesion-region spreading and integrin binding and clustering. In particular, spreading speed may affect cluster number and size by shifting the balance between the formation of new clusters over a wide area and the local growth of pre-existing clusters.

### 4.5 Limitations and future perspectives

A major limitation of the present study is that, although the proposed model reproduced the tendency for clusters to form in the periphery of the adhesion region, it did not sufficiently reproduce the experimentally observed distributions in which almost no clusters were present in the interior. One possible reason for this discrepancy is that the present model does not sufficiently incorporate mechanisms that disassemble integrin clusters formed in the interior of the adhesion region.

The force-dependent dissociation and lateral motion of integrins were represented in simplified forms. Integrin dissociation was modeled as a Bell-type slip bond, assuming that a larger force acting on a bond promotes dissociation. In actual integrin binding, however, catch-bond behavior has been reported, in which bond lifetime increases under low to intermediate tension (Kong et al., 2009; Changede and Sheetz, 2017). If this property is taken into account, immature clusters formed in the interior of the adhesion region, where sufficient tension is not transmitted, may dissociate more readily, whereas clusters along the edge of the adhesion region coupled to the cortical actin layer may be stabilized by an appropriate level of tension. Thus, introducing catch–slip-type force-dependent dissociation may improve the selective reproduction of clustering along the edge of the adhesion region. In addition, integrin lateral mobility was represented using a spatially and temporally constant effective diffusion coefficient. Although its value was selected on the basis of experimentally reported integrin mobility, this approximation does not represent heterogeneity in integrin mobility, cytoskeletal confinement, state-dependent immobilization, or anomalous diffusion in living cell membranes. These simplifications may limit the ability of the model to reproduce the spatial persistence and remodeling of integrin clusters.

Membrane stiffness was treated as temporally constant and spatially uniform in the present model, whereas in real cells the apparent membrane stiffness and membrane tension vary spatiotemporally because of actomyosin contraction, cortex attachment and detachment, and membrane fluctuations (De Belly et al., 2023). In particular, along the edge of the adhesion region, NA formation and maturation are promoted by actin polymerization and actomyosin-derived contractile tension, whereas less stable clusters in the interior may be more readily disassembled by membrane fluctuations and local cortex detachment.

The importance of such mechanochemical coupling has also been suggested by previous computational and experimental studies. A whole-cell mechanochemical migration model coupled integrin dynamics with actin cytoskeletal remodeling through force-dependent integrin–ligand binding and unbinding kinetics, suggesting that adhesion strength, protrusion force, and contractile force jointly regulate cell motion (Feng et al., 2018). In addition, paxillin has been reported to be associated with actomyosin tension and to undergo tension-dependent clustering and turnover (Deguchi et al., 2017). This suggests that the paxillin localization observed along the edge of the adhesion region may correspond to high-tension regions generated by the actomyosin system. Furthermore, a one-dimensional multiscale model of fibroblast–substrate interactions suggested that chemomechanical coupling among focal adhesion dynamics, Ca^2+^ signaling, and stress fiber contractility regulates integrin recruitment, traction force generation, and adhesion stability in a spatially heterogeneous manner (Jaddivada and Gundiah, 2024). However, when interpreting the comparison between the model and experiments in the present study, it is important to note that they represent different molecular observables. The model represents integrin–ligand binding and integrin clustering, whereas the experiments measure paxillin fluorescence, and paxillin recruitment is not included in the model. Consequently, a simulated bound-integrin cluster cannot by itself be identified as a paxillin-positive NA. The simulated clusters should instead be interpreted as putative integrin-based precursors or cores of early adhesion sites, and the comparison with the experimental data is limited to the similarity of their normalized cell-scale spatial patterns and image-derived cluster metrics.

A further limitation concerns the model length scale. Because the simulations used a reduced cell radius, several nondimensional stiffness coefficients that depend on *L*_c_ = *R*_cell_ may be larger than those corresponding to the experimental cell scale, even when the same dimensional material parameters are used. Therefore, when relating the limited spreading under the reference condition and the partial agreement observed under the lower-*λ* conditions to the effective stiffness of the membrane–cortex system in real cells, the influence of the selected model length scale on the nondimensional stiffness coefficients should also be considered.

Future work should therefore account more explicitly for the influence of cell-size scaling on the values of the nondimensional coefficients, while incorporating more detailed representations of force-dependent integrin–ligand binding and unbinding and integrin lateral mobility, together with the spatiotemporal variation in membrane stiffness and membrane tension arising from actomyosin activity, membrane fluctuations, membrane protrusions, and the effects of signaling pathways linking integrins and the cytoskeleton. These extensions would allow the disassembly of interior clusters and the selective stabilization of clustering along the edge of the adhesion region to be examined.

## 5 Conclusion

In this study, we developed a multiscale mechanical model of the early stage of cell adhesion that couples membrane mechanics, integrin transport on the membrane surface, and integrin–ligand binding dynamics. The simulation results showed partial agreement with the experimental observations. In particular, the distribution of bound integrins reproduced the overall profile from the center to the periphery of the adhesion area, the local accumulation near the adhesion edge, and some of the relationships of cluster area, cluster size, and cluster number with the normalized adhesion area. The simulations suggest the following spatial and temporal characteristics of integrin clustering:

- Bound integrins are generally distributed preferentially toward the periphery and can form localized aggregates near the adhesion edge.
- Membrane stiffness and the elapsed time after the initiation of adhesion affect integrin distribution, which may partially explain the cell-to-cell variability observed in the experiments.
- Integrin–ligand binding-induced membrane deformation promotes cluster formation, and its magnitude influences cluster size and spatial distribution.
- The relative timescales of membrane spreading and integrin binding influence the formation and spatial organization of integrin clusters.

More advanced modeling is required to incorporate additional biological effects and may provide a multiscale framework for investigating cell adhesion dynamics across multiple stages of the adhesion process.

## Acknowledgments

The authors thank Yuto Hatsuse and Bansei Andoshiro for their technical assistance. The authors also thank Edanz (https://jp.edanz.com/ac) for editing a draft of this manuscript.

## Statements and Declarations

### Consent for publication

All authors have read and approved the final manuscript and consent to its publication.

### Data availability

The data that support the findings of this study are available from the corresponding author upon reasonable request.

### Author contribution

K.T. contributed to conceptualization, investigation, methodology, software, cell experiments, data analysis, visualization, funding acquisition, and writing original draft.

T.K. contributed to methodology, cell experiments, and data analysis. H.M. and N.S. contributed to methodology, cell experiments, data analysis, review and editing. H.W. contributed to review and editing. S.I. contributed to conceptualization, methodology, software, funding acquisition, supervision, and writing original draft.

