## Supplementary for "Numerical study of spatial and temporal dynamics of integrin clustering during early cell adhesion"

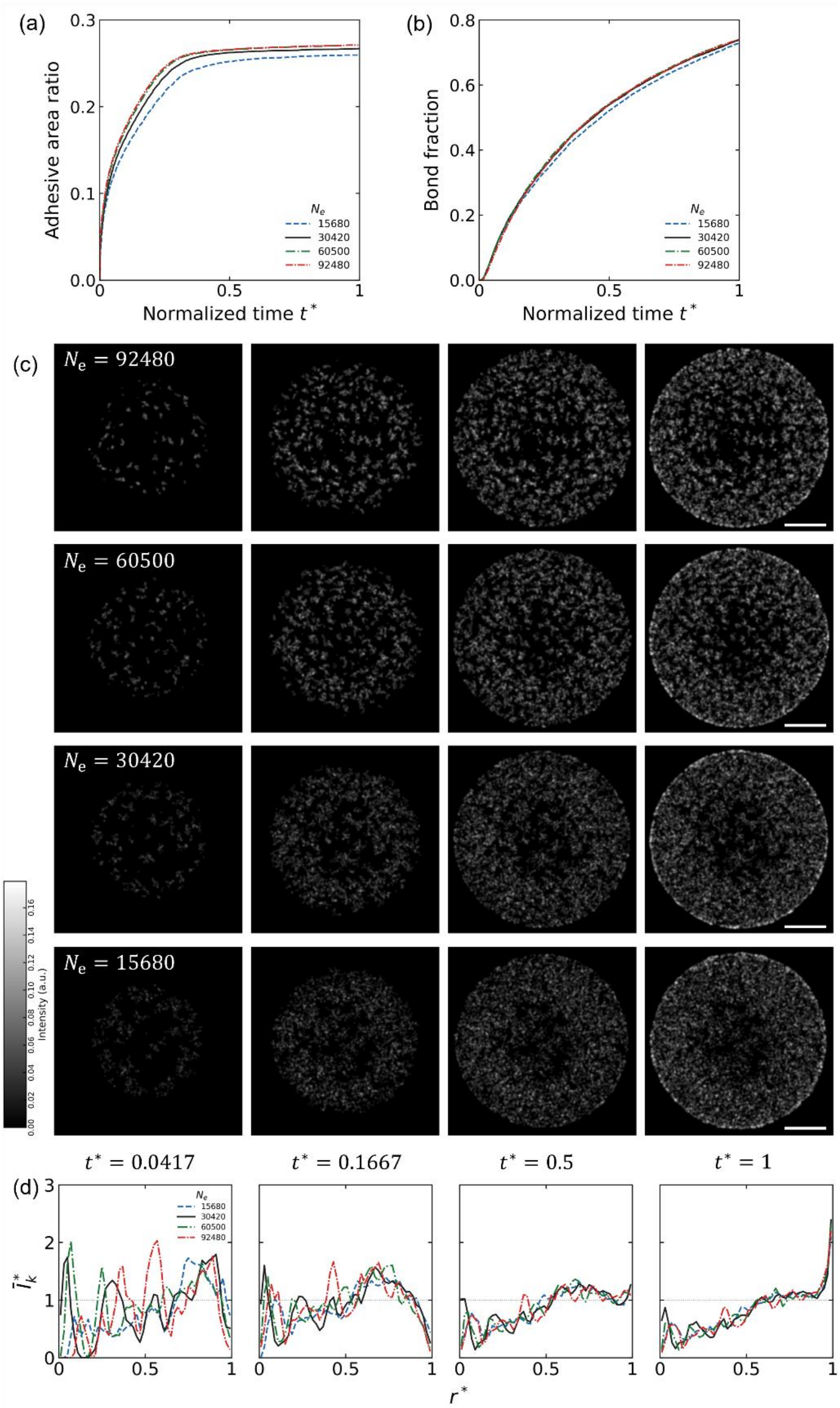


**Fig. S1** Effect of mesh resolution on the simulation results. Time evolution of **a** the normalized adhesion area and **b** the fraction of bound integrins. **c** Integrin pseudo-fluorescence images during cell adhesion obtained using meshes comprising 92,480 elements (top), 60,500 elements (second row), 30,420 elements (third row), and 15,680 elements (bottom). **d** Time evolution of the radial profiles of normalized mean intensity for the corresponding mesh resolutions. The scale bar in **c** represents a normalized length of 0.5.

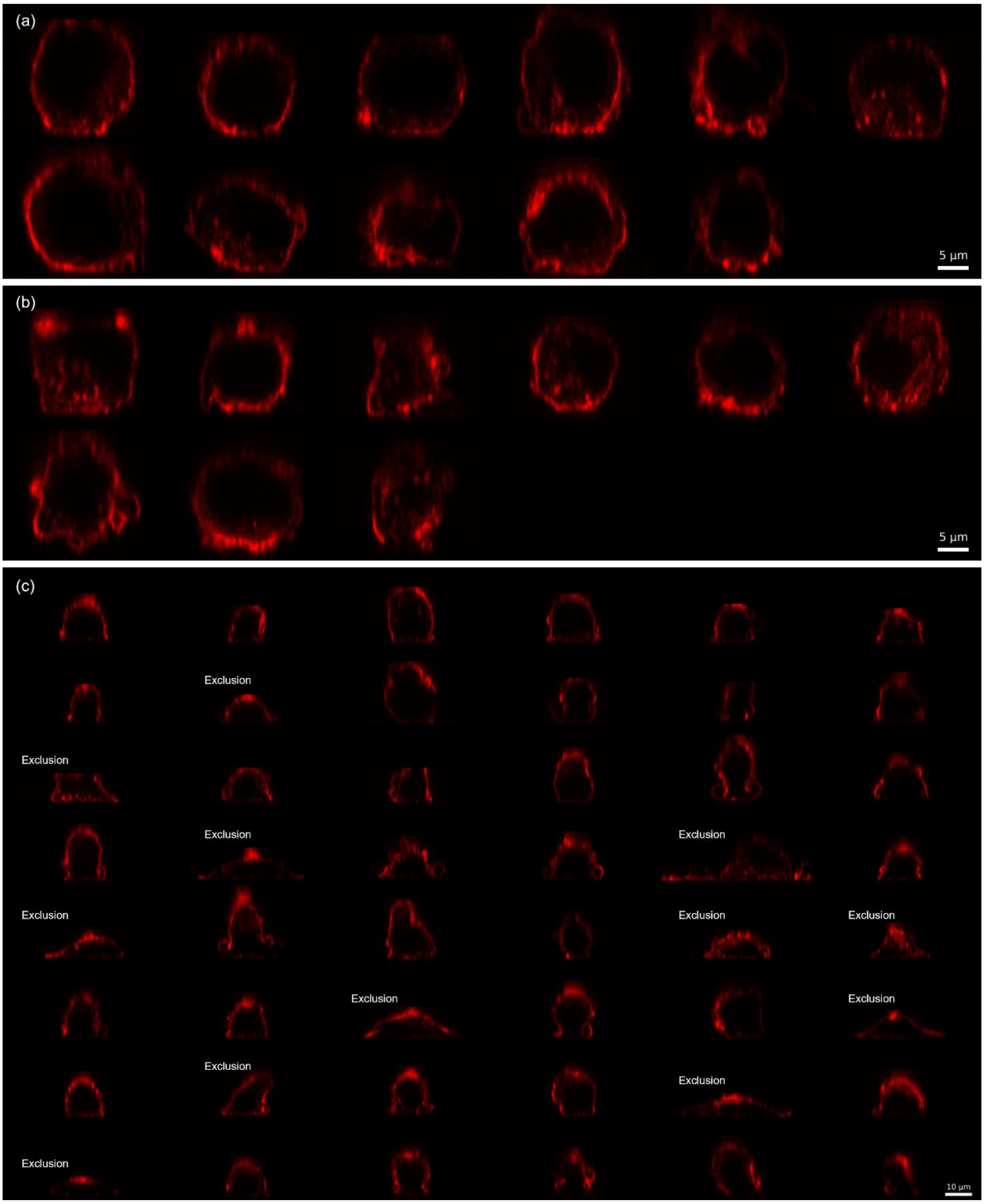


**Fig. S2** Lateral (XZ) fluorescence images of actin at 2 min (**a**), 5 min (**b**), and 15 min (**c**). Scale bars represent 5 μm in (**a**, **b**) and 10 μm in (**c**). In (**c**), cells excluded from the quantitative analysis are labeled "Exclusion." Twelve cells showing advanced spreading morphology, including marked flattening, pronounced lamellipodia, or multiple prominent filopodia, were excluded.

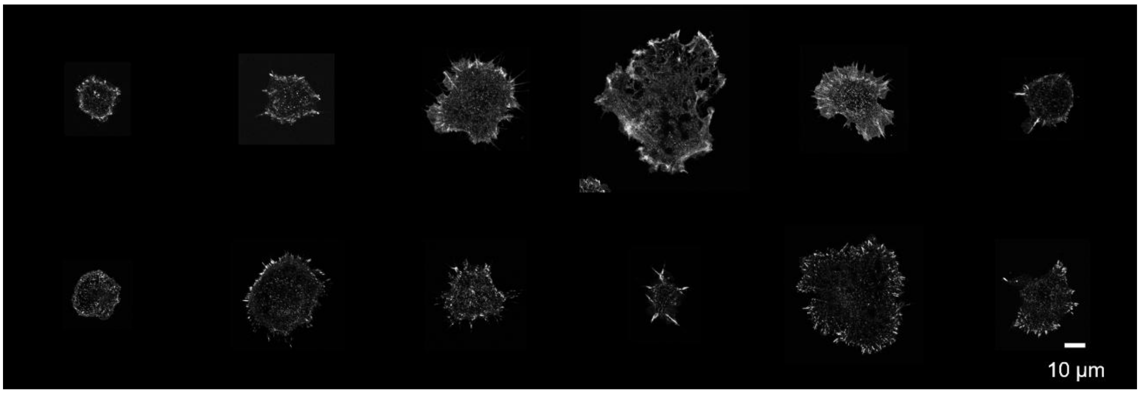


**Fig. S3** Paxillin distribution of excluded samples (n = 12). The scale bar indicates 10 μm. Twelve cells showing advanced spreading morphology, including marked flattening, pronounced lamellipodia, or multiple prominent filopodia, were excluded.

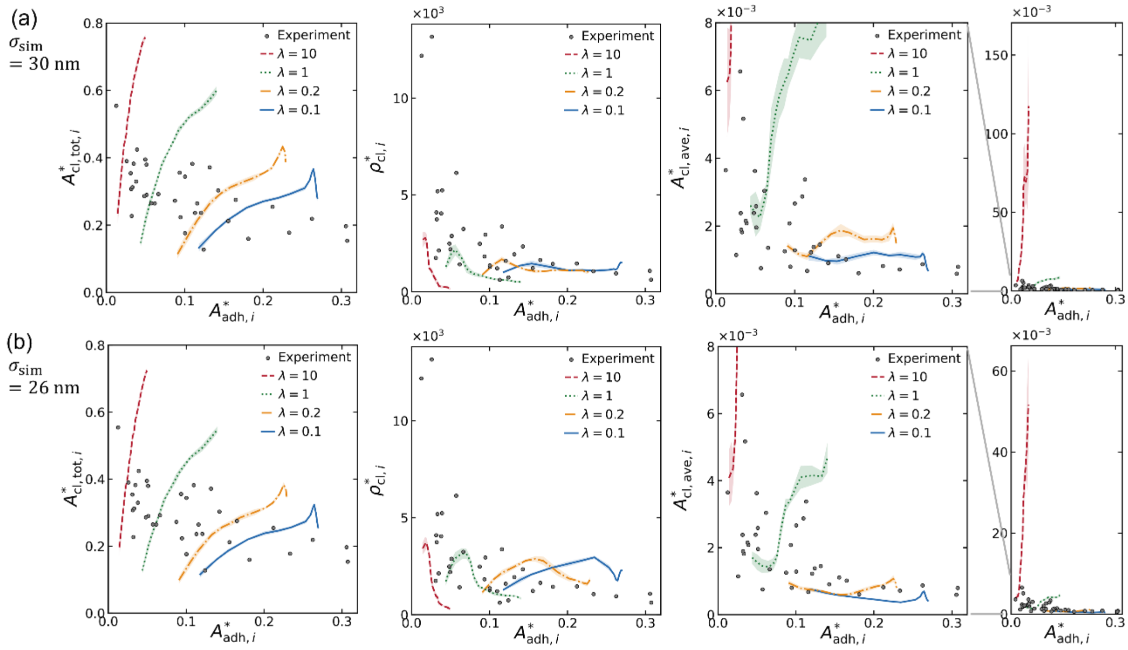


**Fig. S4** Effect of the Gaussian point-spread-function width, $\sigma_{\text{sim}}$, on cluster analysis. Results obtained using **a** $\sigma_{\text{sim}} = 30$ nm and **b** $\sigma_{\text{sim}} = 26$ nm. The horizontal axis represents the normalized adhesion area $A^*_{\text{adh},i}$; the vertical axes show the total cluster area fraction $A^*_{\text{cl,tot},i}$ (left), normalized cluster number density $\rho^*_{\text{cl},i}$ (center), and normalized mean cluster area $A^*_{\text{cl,ave},i}$ (right).

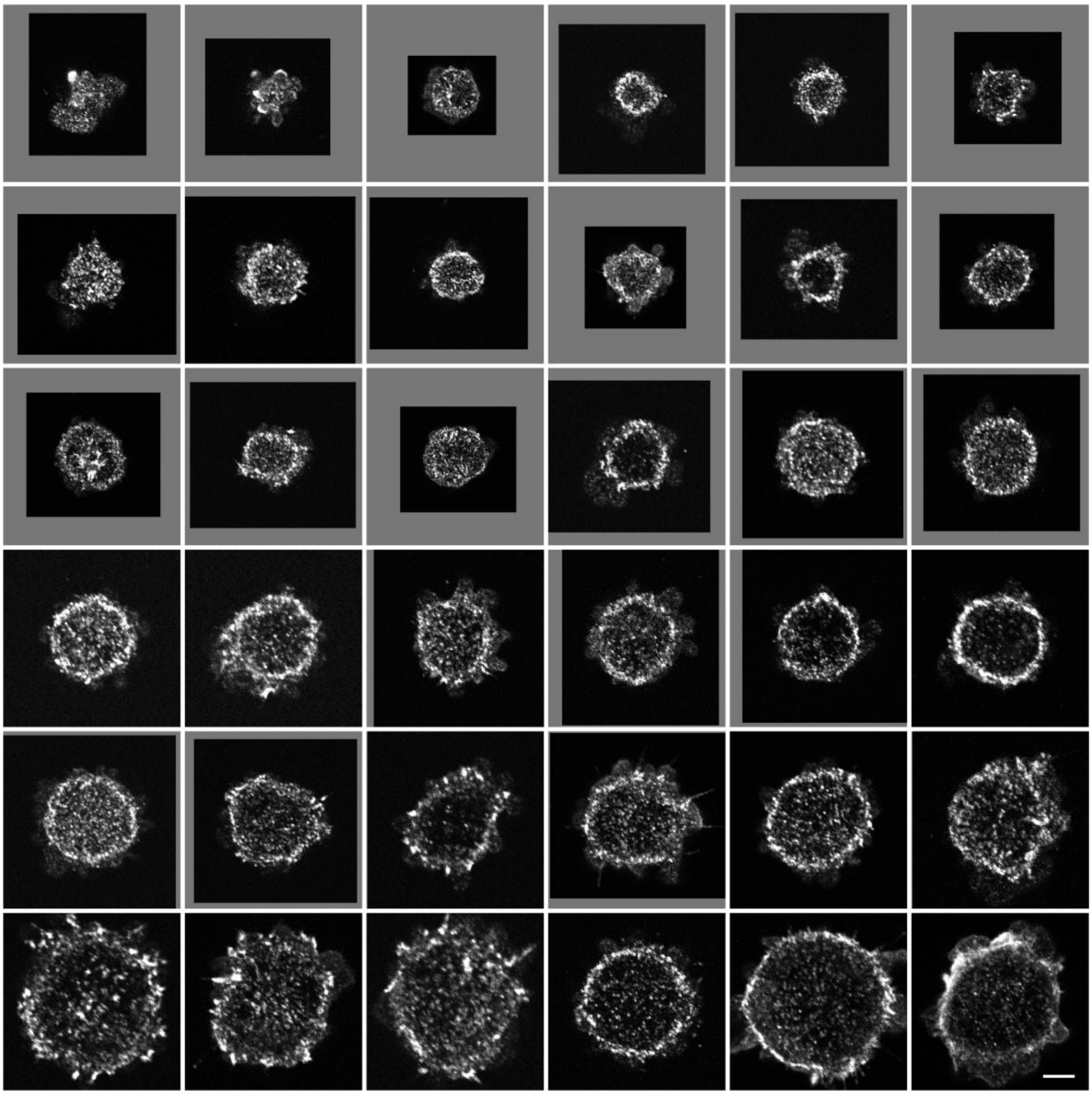

**Fig. S5** Paxillin distribution (n = 36). The image in the $k$-th row and $l$-th column represents a cell belonging to the $k$-th sextile of $A^*_{\mathrm{adh}}$ and having the $l$-th smallest $Q_{\mathrm{periphery}}$ within that sextile. The scale bar indicates a normalized length of 0.5.

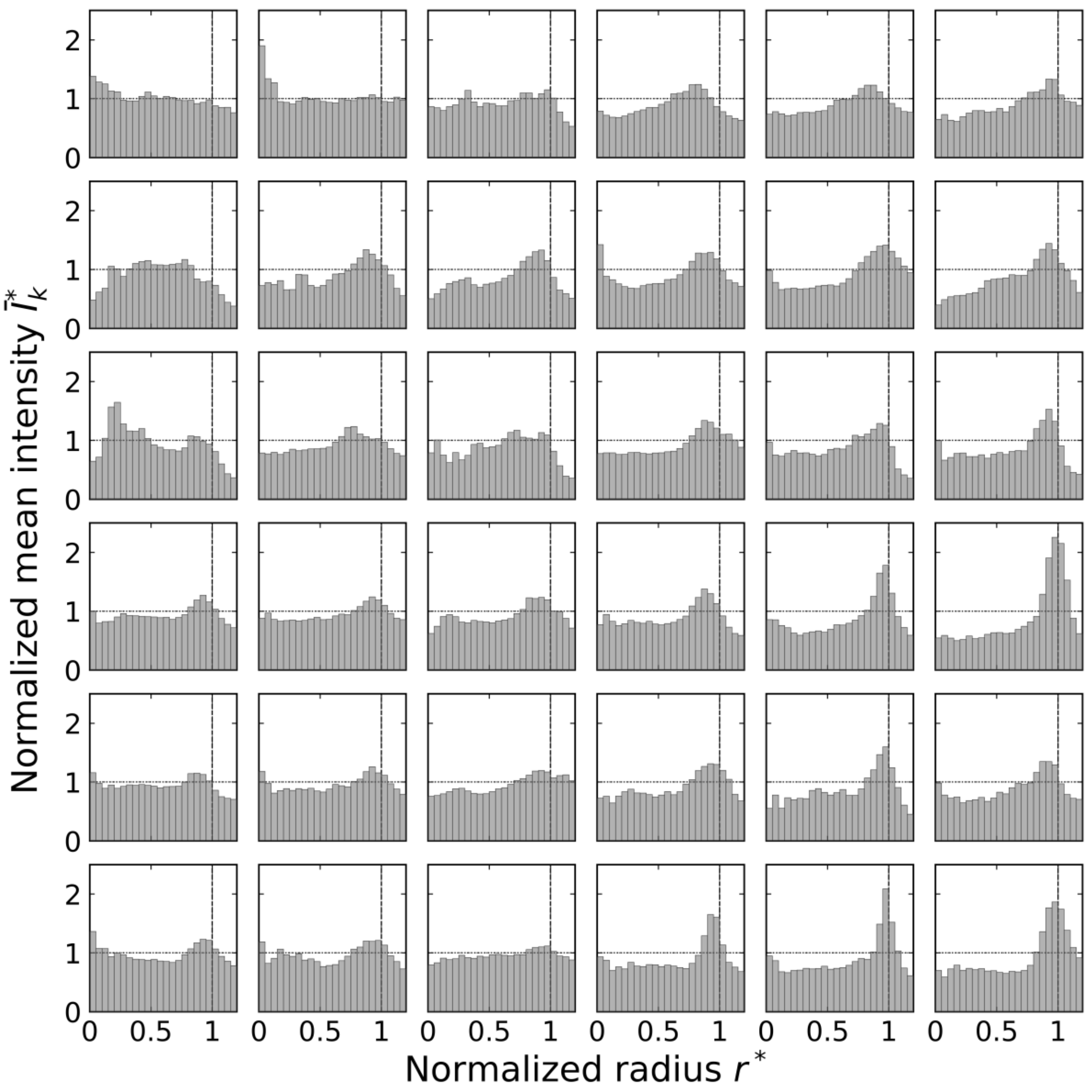


**Fig. S6** Radial distributions of normalized mean paxillin fluorescence intensity ($n = 36$). The plot in the $k$-th row and $l$-th column represents a cell belonging to the $k$-th sextile of $A^*_{\mathrm{adh}}$ and having the $l$-th smallest $Q_{\mathrm{periphery}}$ within that sextile. Each bar represents the normalized mean intensity within an annular region of normalized radial width $\Delta r^* = 0.05$. The displayed range of normalized mean intensity was fixed at $[0,1.2]$ for all panels.

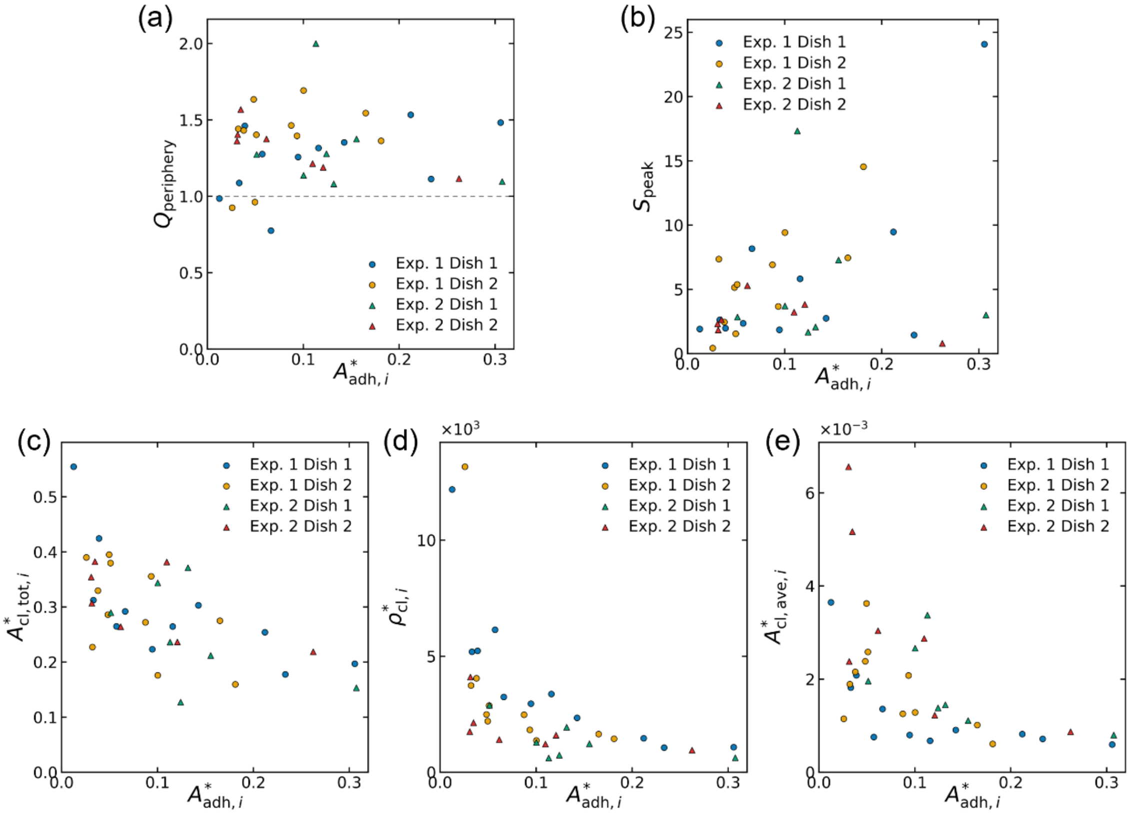


**Fig. S7** Relationships between the normalized adhesion area and spatial-distribution and cluster metrics in the experiments. Relationships of the normalized adhesion area $A^*_{\mathrm{adh},i}$ with **a** the peripheral-to-central mean intensity ratio $Q_{\mathrm{periphery}}$, **b** the peripheral peak sharpness $S_{\mathrm{peak}}$, **c** the total cluster area fraction $A^*_{\mathrm{cl,tot},i}$, **d** the normalized cluster number density $\rho^*_{\mathrm{cl},i}$, and **e** the normalized mean cluster area $A^*_{\mathrm{cl,ave},i}$. Each point represents one of the 36 cells examined at $T_{\mathrm{inc}} =$ 15 min. The cells were obtained from four dishes in two independent experiments (7–11 cells per dish), and the four experiment–dish groups are identified in the legends. The dashed line in **a** indicates $Q_{\mathrm{periphery}} = 1$, corresponding to equal mean intensities in the peripheral and central regions.

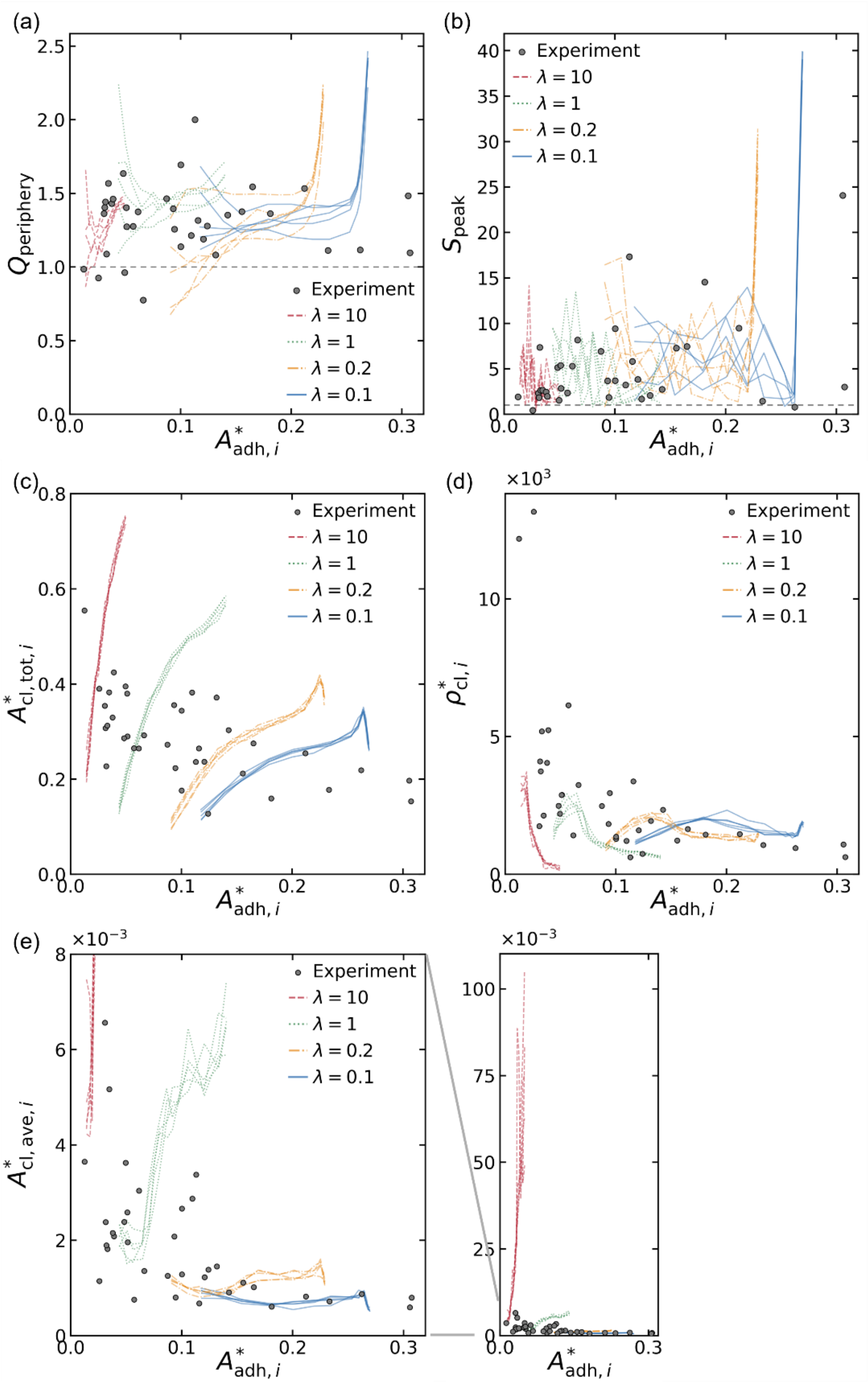


**Fig. S8** Individual simulation trajectories for each membrane-stiffness condition, obtained using five different random seeds. Each trajectory represents one random-seed realization. The horizontal axis in each panel represents the normalized adhesion area $A^*_{\mathrm{adh},i}$; the vertical axes show **a** the peripheral-to-central mean intensity ratio $Q_{\mathrm{periphery}}$, **b** the peripheral peak sharpness $S_{\mathrm{peak}}$, **c** the total cluster area fraction $A^*_{\mathrm{cl,tot},i}$, **d** the normalized cluster number density $\rho^*_{\mathrm{cl},i}$, and **e** the normalized mean cluster area $A^*_{\mathrm{cl,ave},i}$.
